# The maize *Hairy Sheath Frayed1* (*Hsf1*) mutant alters leaf patterning through increased cytokinin signaling

**DOI:** 10.1101/743898

**Authors:** Michael G. Muszynski, Lindsay Moss-Taylor, Sivanandan Chudalayandi, James Cahill, Angel R. Del Valle-Echevarria, Ignacio Alvarez-Castro, Abby Petefish, Hitoshi Sakakibara, Dmitry M. Krivosheev, Sergey N. Lomin, Georgy A. Romanov, Subbiah Thamotharan, Bailin Li, Norbert Brugière

## Abstract

Leaf morphogenesis requires growth polarized along three axes - proximal-distal, medial-lateral and abaxial-adaxial. Grass leaves display a prominent proximal-distal (P-D) polarity consisting of a proximal sheath separated from the distal blade by the auricle and ligule. Although proper specification of the four segments is essential for normal morphology, our knowledge is incomplete regarding the mechanisms which influence P-D specification in monocots like maize (*Zea mays*). Here we report the identification of the gene underlying the semi-dominant, leaf patterning, maize mutant *Hairy Sheath Frayed1* (*Hsf1*). *Hsf1* plants produce leaves with outgrowths consisting of proximal segments – sheath, auricle and ligule – emanating from the distal blade margin. Analysis of three independent *Hsf1* alleles revealed gain-of-function missense mutations in the ligand binding domain of the maize cytokinin (CK) receptor *Zea mays Histidine Kinase1* (*ZmHK1)* gene. Biochemical analysis and structural modeling suggest the mutated residues near the CK binding pocket affect CK binding affinity. Treatment of wild type seedlings with exogenous CK phenocopied the *Hsf1* leaf phenotypes. Results from expression and epistatic analyses indicated the *Hsf1* mutant receptor appears to be hypersignaling. Our results demonstrate that hypersignaling of CK in incipient leaf primordia can reprogram developmental patterns in maize.

**Summary:** Increased cytokinin signaling in the maize *Hairy Sheath Frayed1* mutant modifies leaf development leading to changes in pattering, growth and cell identity.

## INTRODUCTION

Proper leaf morphogenesis in higher plants requires defined patterns of growth polarized along three axes: adaxial-abaxial, medial-lateral and proximal-distal (McConnell and Barton, 1998; Tsukaya, 1998; Bowman *et al*., 2002; Byrne *et al*., 2002). Growth along the proximal-distal (P-D) axis is particularly evident in grass leaves, like maize, which are composed of four distinct segments; the sheath is proximal, the blade is distal and the auricle and ligule form the boundary between the two (Figure 1A) (Sylvester *et al*., 1996). A number of genes have been identified that influence P-D patterning, with *BLADE-ON-PETIOLE* (*BOP*) genes affecting proximal identity in eudicots and monocots (Ha *et al*., 2003, 2004; Norberg *et al*., 2005; Toriba *et al*., 2019; Moon *et al*., 2013; Tavakol *et al*., 2015). In grasses, ectopic expression of class I *knotted1like homeobox* (*knox*) transcription factor genes in developing leaf primordia alters P-D patterning, primarily disrupting the formation of a defined sheath-blade boundary (Freeling and Hake, 1985; Hake *et al*., 1989, 1991; Smith *et al*., 1992; Schneeberger *et al*., 1995; Muehlbauer *et al*., 1997; Foster *et al*., 1999a; Tsiantis *et al*., 1999; Byrne *et al*., 2001). Class I *knox* genes typically function in meristem formation and maintenance, and their down-regulation is required for normal development of determinant organs like leaves (Endrizzi *et al*., 1996; Long *et al*., 1996; Kerstetter *et al*., 1994). In meristems, KNOX proteins function to increase cytokinin (CK) accumulation by positive regulation of CK synthesis genes and simultaneously decrease gibberellic acid (GA) accumulation by suppression of GA biosynthesis genes or activation of GA catabolic genes (Ori et al., 2000; Sakamoto et al., 2001; Hay et al., 2002; Jasinski et al., 2005; Yanai et al., 2005; Sakamoto et al., 2006; Bolduc and Hake, 2009). In addition, a rice KNOX transcription factor was shown to also affect brassinosteroid (BR) accumulation by upregulating BR catabolism in the shoot apical meristem (Tsuda *et al*., 2014). Determinate leaf primordia form when *knox* expression is down-regulated by the action of ROUGH SHEATH2 (RS2) and related proteins resulting in a decrease in CK and increase in GA accumulation (Hay *et al*., 2006; Guo *et al*., 2008). In addition to the action of CK and GA, auxin is required for proper leaf initiation and positioning. The polar transport of auxin by PINFORMED1 (PIN1) auxin efflux carriers guides the formation of auxin maxima, localized regions of high auxin accumulation, that is required for initiation of leaf primordia (Pozzi *et al*., 2001; Scarpella *et al*., 2006; Benjamins and Scheres, 2008; Zhao, 2008). The emerging model predicts that spatial differences in cytokinin/auxin ratios control final cell fate (Shani *et al*., 2006; Muller and Sheen, 2008). Ectopic *knox* expression presumably shifts critical phytohormone ratios in developing leaf primordia but the exact molecular mechanisms by which phytohormone ratios determine leaf patterning remain incomplete.

**Figure 1.**
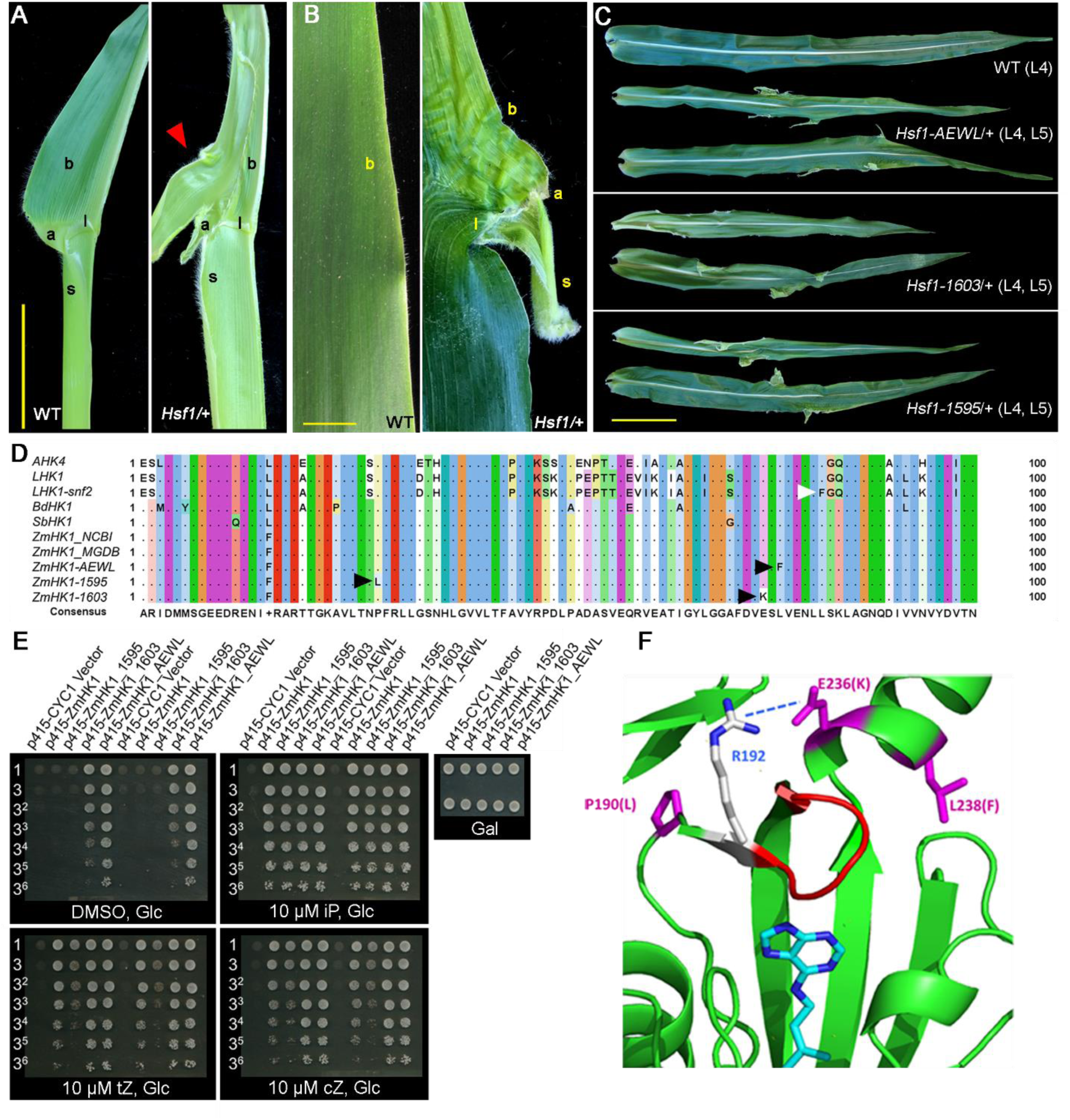
*Hsf1* mutants alter leaf patterning and are caused by missense mutations in the *ZmHK1* cytokinin receptor. **(A)** Adaxial view of half-leaves from WT and *Hsf1-1603/+* sibs showing the proximal-distal organization of the sheath (s), ligule (l), auricle (a) and blade (b) and a prong outgrowth (red triangle). Bar = 5 cm. **(B)** Close-up of a blade margin (b) from WT and *Hsf1-1603/+* showing a prong consisting of proximal leaf segments – sheath (s), ligule (l) and auricle (a) juxtaposed to the blade (b). Bar = 1 cm. **(C)** Comparison of leaf phenotypes between the three *Hsf1* alleles. L4 (top), 4^th^ leaf below tassel; L5 (bottom, 5^th^ leaf below tassel. Bar = 10 cm. **(D)** Amino acid alignment of a portion of the CHASE domain from different plant his-kinase cytokinin receptors and the three *Hsf1* mutant alleles. Missense residues are marked by black triangles for the *Hsf1* alleles and by a white triangle for the *Lotus snf2* allele. Amino acid sequences derived from AT2G01830 (*AHK4*), AM287033 (*LHK1* and *LHK1-snf2*), XM_003570636 (*BdHK1*), XM_002454271 (*SbHK1*), NM_001111389 (*ZmHK1-NCBI*), GRMZM2G151223 (*ZmHK1-MaizeGDB*), *ZmHK1* from the A619 inbred (*ZmHK1-AEWL*) and the Mo17 inbred (*ZmHK1-1603* and *ZmHK1-1595*). **(E)** ZmHK1 receptors with *Hsf1* mutations show CK independent growth in a yeast his-kinase signaling assay. Growth of *S. cerevisiae sln* Δ mutant transformed with an empty vector, the ZmHK1 vector or one of the *Hsf1* mutant *ZmHK1* vectors on glucose media with no CK (DMSO) or supplemented with different cytokinins - iP, tZ, or cZ. Growth on galactose media of the *sln* Δ mutant transformed with each of the assayed vectors. DMSO, dimethyl sulfoxide; iP, *N^6^*-(*Δ^2^*-isopentenyl)adenine; tZ, *trans*-zeatin; cZ, *cis*-zeatin. Dilutions of yeast cultures (O.D._600_ = 1.0) for each yeast strain are noted on the left of each image. (F) Ribbon diagram of the ZmHK1 CHASE domain with the *Hsf1* mutations (magenta) noted and one molecule of *N^6^*-(*Δ^2^*-isopentenyl)adenine (blue and aqua) complexed in the binding pocket. Arginine 192 (blue), in the loop domain (red) forming one face of the binding cavity, is predicted to form a salt bridge with E236, the residue altered in *Hsf1-1603*. *Hsf1-1595* is P190L, *Hsf1-1603* is E236K and *Hsf1-AEWL* is L238F.

As phytohormones play pivotal roles in many developmental programs, the pathways that signal their perception and response have been well characterized. For example, the perception and response to the CK phytohormones relies on a two-component signal transduction system (Hwang and Sheen, 2001; Yonekura-Sakakibara *et al*., 2004; Hwang and Sakakibara, 2006; Du *et al*., 2007; To and Kieber, 2008). The perception of CK is mediated via a partially redundant signaling system of histidine kinases (HKs), histidine phosphotransfer proteins (HPTs) and response regulators (RRs). CK signaling begins with the perception of CK by binding to HK receptors at the ER, and probably also plasma membrane, which triggers receptor phosphorylation (Lomin *et al*., 2011). The activated receptors initiate phosphorelay by transferring phosphoryl groups to HPTs, which shuttle between the cytoplasm and nucleus. Once in the nucleus, phosphorylated HPTs transfer their phosphoryl groups to type-B RRs, which in turn activate expression of type-A RRs and other CK responsive genes (Rashotte *et al*., 2006). The type-A RRs and other CK-responsive genes mediate several CK-regulated processes including shoot and root growth, de-etiolation, leaf expansion, root vascular development, senescence, and cytokinin homeostasis (To and Kieber, 2008). In maize, multiple members of each of the CK signaling components have been identified (Yonekura-Sakakibara *et al*., 2004). Maize has seven HKs (ZmHKs), of which, three have been shown to bind and signal various types of CKs in heterologous assays (Lomin *et al*., 2011; Steklov *et al*., 2013). Three HPTs (ZmHPs), three type-B RRs and seven type-A RRs (ZmRRs) have also been identified in maize (Asakura *et al*., 2003). Of these signal transduction components, the function of only *ZmRR3*, a type-A RR, has been defined by null mutations and shown to underlie the *aberrant phyllotaxy1* (*abph1*) mutation (Jackson and Hake, 1999; Giulini *et al*., 2004). Our understanding of the functions of other components of the CK signal transduction pathway remains incomplete for cereal species like maize.

To gain a better understanding of the signaling mechanisms which mediate leaf pattern specification, we initiated a study of the semi-dominant *Hairy Sheath Frayed1* (*Hsf1*) mutation which alters P-D leaf development in maize (Bertrand-Garcia and Freeling, 1991a). Although *Hsf1* disrupts the P-D leaf pattern similar to dominant class I *knox* mutations, *Hsf1* is not itself a *knox* gene, since it does not map to the location of any maize *knox* genes (Bertrand-Garcia and Freeling, 1991b). In this report, we show that the *Hsf1* phenotype results from specific missense mutations in the maize CK receptor *Zea mays Histidine Kinase1* (*ZmHK1*) gene (Yonekura-Sakakibara *et al*., 2004). Our analyses of mutant receptor function, the effects of exogenous CK treatment on leaf development, and epistatic interaction suggest that the ZmHK1 receptor is hypersignaling in *Hsf1* mutants. Overall, our results indicate CK hypersignaling can influence the specification of P-D leaf patterning in maize and underscores the capacity of CK to alter developmental programs.

## RESULTS

### The *Hsf1* mutation induces specific alterations to maize leaf patterning

The original *Hsf1* mutation arose via ethyl methanesulfonate (EMS) mutagenesis of the inbred line Mo17 and was designated *Hsf1-N1595* (also called *Hsf1-O*) (Bird and Neuffer, 1985). A second mutation, *Hsf1-N1603* (hereafter called *Hsf1-1603*), was shown to be allelic or very closely linked (Bertrand-Garcia and Freeling, 1991a). We isolated three additional alleles in independent EMS mutagenesis screens in different inbred backgrounds: *Hsf1-AEWL* in A619, *Hsf1-2559* in Mo17, and *Hsf1-7322* in A632. All *Hsf1* alleles have very similar phenotypes compared to the *Hsf1-N1595* (hereafter called *Hsf1-1595*) reference mutation. As was shown previously for *Hsf1-1595*, plants heterozygous for any of the *Hsf1* alleles display a highly penetrant mutant leaf patterning phenotype with outgrowths consisting of sheath, auricle and ligule emanating from the distal blade margin (Figures 1A to 1C) (Bertrand-Garcia and Freeling, 1991a). The outgrowths have proximal identity and were termed “prongs”, which we adopted to describe this structure (Figure 1B). Although *Hsf1* mutant plants have proximal tissue growing on the distal blade, they have a normal blade-sheath boundary (Figure 1A) (Bertrand-Garcia and Freeling, 1991a). All the pleiotropic phenotypes described for *Hsf1-1595* in Bertrand-Garcia and Freeling (1991a) are shared by all the other *Hsf1* alleles, including an increase in macrohair size and density on the abaxial sheath, adaxial blade, and blade margin, an increase in leaf number, shorter stature, short and narrow leaves, and reduced root growth (Supplemental Figures 1A to 1B; Supplemental Table 1). Bertrand-Garcia and Freeling (1991a) also showed homozygous *Hsf1* plants have a stronger mutant phenotype being extremely stunted, with multiple shoots arising from the coleoptile node at germination, and having adventitious needle- or club-shaped leaves (Supplemental Figures 1A to 1B).

Since plants heterozygous for ether *Hsf1-1595*, *Hsf1-1603*, or *Hsf1-AEWL* were phenotypically very similar (Figure 1C), we chose the *Hsf1-1603* allele to characterize the temporal and spatial patterns of prong formation to better understand how the *Hsf1* mutation affected leaf patterning. In *Hsf1-1603* heterozygotes, prongs first appeared on leaf 5 in a few plants, and most commonly appeared on leaf 6 or leaf 7 but never on earlier arising leaves (Supplemental Figure 1C). The earliest sign of P-D leaf polarity specification is the formation of the preligule band (PLB) which will differentiate into the auricle and ligule (Sylvester *et al*., 1990; Johnston *et al*., 2014). Formation of the PLB typically is first observed in plastochron 5 or 6 stage leaf primordia (P5 - P6) with the initiating ligule becoming visible about plastochron 7 or 8 (P7 - P8) (Johnston *et al*., 2014). Plastochron describes the stage of leaf primordia development and refers to the position of the primordia relative to the shoot apical meristem (SAM) (Sylvester *et al*., 1990). Thus, a P5 primordium has four younger primordia between it and the SAM. To determine if the *Hsf1-1603* mutation influenced the timing of the acquisition of P-D polarity, we examined leaf primordia in *Hsf1-1603/+* and wild type sib plants for signs of early ligule development (see Methods). The initiating ligule was most commonly first visible on P7 primordia in both wild type and *Hsf1-1603* heterozygotes indicating no influence on P-D polarity acquisition (Supplemental Figure 1E). To determine if the appearance of prong primordia on the blade margin coincided with the acquisition of P-D polarity, developing leaf primordia from *Hsf1-1603* heterozygotes were dissected and examined for the presence of initiating prongs. Prong initials were most commonly observed on the blade margins of P5 or P6 leaf primordia but some were noted as early as P4 (Supplemental Figure 1D, 1F to 1G), consistent with prong formation in *Hsf1-1595* heterozygotes (Bertrand-Garcia and Freeling, 1991b). Thus prongs typically initiated from blade margins about the same plastochron stage as formation of the PLB.

Prongs were observed to occur in different sizes and at different positions along the leaf blade margin (Figures 1B and 1F, and Supplemental Figure 2A). To determine if prong formation was random or patterned, we measured the size and positon of each prong from both margins of mature leaves collected from different positions on the shoot of *Hsf1-1595*, *Hsf1-1603* and *Hsf1-AEWL* heterozygous plants. Results showed that prong formation was more frequent on leaves higher on the shoot (Supplemental Table 2) with prongs occupying more of the blade margin in these upper leaves compared to lower leaves (Supplemental Figures 2B and 2C). Next we determined where prongs formed along the P-D axis of the blade. Analysis indicated prongs only formed in the proximal 70% and never in the distal 30% of the blade, with the majority of prongs forming within a region encompassing the proximal 15% to 40% of the blade (Supplemental Figure 2D). Next we examined the range of prong sizes for each *Hsf1* allele within this prong-forming region. For all three alleles, the majority of prongs were about a centimeter in size but a few were larger, ranging from 3 to 6 centimeters (Supplemental Figure 2E). With relative position and size known, we next asked whether prong position was related to its size. In general, the largest prongs often formed in the basal 20% of the blade and smaller prongs formed at any position within the prong forming region (Supplemental Figure 2F). Thus, our analysis indicated prong formation was not random but occurred in particular regions of the blade and initiated at specific developmental stages.

### Gain-of-function mutations in the maize cytokinin receptor gene *ZmHK1* underlie the *Hsf1* mutation

Previous studies mapped *Hsf1-1595* to the long arm of chromosome 5 (Bertrand-Garcia and Freeling, 1991b). To isolate the gene underlying the *Hsf1* locus, we screened a backcross mapping population of over 3,000 plants with linked molecular markers derived from the maize reference genome (B73 RefGen_v1). The *Hsf1* locus was localized to a 21-kb interval with a single gene model (GRMZM2G151223, B73 RefGen_v2). This gene model was well supported with abundant EST evidence and was annotated as encoding *Zea mays Histidine Kinase1* (*ZmHK1*), one of seven maize histidine kinase cytokinin receptors (Yonekura-Sakakibara *et al*., 2004; Steklov *et al*., 2013). To confirm *ZmHK1* was the correct gene and to identify the causative lesions, the *ZmHK1* gene was sequenced from all five *Hsf1* alleles. The entire *ZmHK1* genomic region, including ca. 2-kb upstream and downstream of the transcription start and stop, was sequenced from *Hsf1-1595*, *Hsf1-1603, Hsf1-2559, Hsf1-7322* and *Hsf1-AEWL* homozygotes and their progenitor inbred lines. As expected for EMS-generated mutations, single nucleotide transitions were identified in the five *Hsf1* alleles compared to their progenitor sequences. Although each allele arose independently, *Hsf1-1595* and *Hsf1-1603* had the exact same transition mutations as *Hsf1-7322* and *Hsf1-2559*, respectively. Thus, hereafter, we refer to the three different *Hsf1* alleles: *Hsf1-1595*, *Hsf1-1603* and *Hsf1-AEWL*. Each transition mutation produced a missense mutation in a highly conserved amino acid located in the CHASE (<u>c</u>yclases/<u>h</u>istidine-kinase-<u>a</u>ssociated <u>se</u>nsory) domain of the ZmHK1 protein, where CK binding occurs (Figure 1D) (Hothorn *et al*., 2011; Steklov *et al*., 2013). The *Hsf1-1595* mutation changed proline 190 to leucine (CCA>CTA), the *Hsf1-1603* mutation changed glutamate 236 to lysine (GAG>AAG), and the *Hsf1-AEWL* mutation changed leucine 238 to phenylalanine (CTT>TTT). The missense mutation in *Hsf1-AEWL* is particularly significant because this is the same type of amino acid substitution, although at a slightly different position in the CHASE domain, which was found in another gain-of-function mutation in a CK receptor, the *spontaneous nodule formation2* (*snf2)* mutation in the lotus *Lhk1* receptor, (Figure 1D) (Tirichine *et al*., 2007). The *snf2* mutation was shown to cause mutant LHK1 to signal independent of the CK ligand in a heterologous signaling assay suggesting the *snf2* mutation locked LHK1 in an active signaling state (Tirichine *et al*., 2007). Based on the location and nature of the amino acid substitutions in the three *Hsf1* mutations and the presumed mode of action of the *snf2* mutation in *Lhk1*, we hypothesized that the *Hsf1* mutations might also lock the ZmHK1 receptor in an active CK signaling state and signal independent of CK.

### The *Hsf1* mutant CK receptors have altered histidine kinase signaling and ligand binding activities

To determine if the *Hsf1* mutant receptors are signaling independent of CK, we utilized a heterologous histidine kinase signaling assay system developed in the yeast *Saccharomyces cerevisae* (Suzuki *et al*., 2001). In the yeast assay, the cognate his-kinase of an endogenous two-component phosphorelay signal transduction system was deleted. Functional replacement of the endogenous his-kinase with the assayed CK receptor, in this case ZmHK1, allowed the activity of the receptor to be determined as the output of the endogenous yeast transduction system, which is the ability to grow on glucose media (Suzuki *et al*., 2001). We engineered the exact point mutation found in each *Hsf1* mutation into the *ZmHK1* cDNA in the p415CYC-ZmHK1 plasmid for expression in yeast (Suzuki *et al*., 2002; Higuchi *et al*., 2009). We next tested receptor activity in the *sln1* deletion yeast strain TM182 carrying each of the *Hsf1* missense mutations, the wild type *ZmHK1* cDNA, and the empty p415CYC vector grown on glucose media with and without the CK ligand (Figure 1E). As expected, the wild type ZmHK1 strain only grew well on glucose media supplemented with higher concentrations of the three CKs tested (Figure 1E) and, at lower CK concentrations, only grew robustly on glucose with the preferred ligand *N^6^*-(*Δ^2^*-isopentenyl) adenine (iP) (Supplemental Figures 3A to 3C). In the absence of added CK, strains carrying either ZmHK1-AEWL or ZmHK1-1603 grew robustly on glucose media (Figure 1E). This result indicated that the ZmHK1-AEWL and ZmHK1-1603 receptors signaled independent of added CK in this assay. To determine if the mutant receptors were still CK responsive, they were also grown on glucose media supplemented with the three tested CKs (Figure 1E). Growth on glucose supplemented with different CKs did not reveal any receptor activity differences between ZmHK1-AEWL and ZmHK1-1603. Surprisingly, growth of the ZmHK1-1595 strain was different than the other two mutant receptors and wild type. The ZmHK1-1595 strain did not grow on glucose media without added CK, similar to wild type ZmHK1 (Figure 1E). Instead, the ZmHK1-1595 strain showed strong growth on glucose media with 10 µM of the preferred CK iP and weaker growth on glucose with 10 µM of two other bioactive CKs, *trans*-zeatin (tZ) and *cis*-zeatin (cZ), suggesting ZmHK1-1595 had weak receptor activity in this assay (Figure 1E

To investigate ligand specificity differences, CK ligand binding affinities were determined for the mutant and wild type receptors (Romanov *et al*., 2005; Lomin *et al*., 2011). Affinities were determined for 6 different CKs and adenine (Ade) using two binding assays with receptors expressed in bacterial spheroplasts (Romanov *et al*., 2005) or residing in tobacco membranes after transient expression *in planta* (Lomin *et al*., 2015, 2011). The ligand preferences for the wild type ZmHK1 receptor were comparable to those determined previously (Table 1) (Lomin *et al*., 2015, 2011). The mutant receptors, on the other hand, all showed increased affinities for most of the CKs tested (Table 1). The preference ranking of the mutant receptors for different CKs was mostly similar to wild type (Supplemental Figure 4) but the affinities were increased between 2- to 8-fold (Table 2). The only exception was the affinity for the synthetic CK thidiazuron, which was reduced for all the mutant receptors compared to wild type ZmHK1. Thus, the missense mutations in the *Hsf1* alleles increased the relative binding affinity of the receptor for all the natural CKs tested, suggesting the mutant receptors might be hypersignaling.

**Table 1.** Apparent affinity constants KD* for wild type and mutant ZmHK1 receptors with different cytokinins.

| $K_D^*$ for cytokinins (nM) | | | | | | | | | |
| --- | --- | --- | --- | --- | --- | --- | --- | --- | --- |
| Assay | Receptor | iP | BA | tZ | cZ | Kin | TD | DZ | Ade |
| Bacterial spheroplasts | ZmHK1 | 2.90 | 3.69 | 31.8 | 37.5 | 33.0 | 37.6 | 312.0 | >10000 |
|  | *AEWL* | 0.36 | 0.56 | 6.38 | 5.56 | 7.62 | 93.7 | 61.6 | >10000 |
|  | *1603* | 0.59 | 0.91 | 7.27 | 6.74 | 7.50 | 111.0 | 88.0 | >10000 |
| Tobacco membrane | ZmHK1 | 0.52 | 1.42 | 7.16 | 8.31 | - | 49.2 | 114.0 | >10000 |
|  | *1595* | 0.23 | 0.31 | 1.65 | 2.14 | - | 71.9 | 14.1 | >10000 |

iP, $N^6$ -( $\Delta^2$ -isopentenyl)adenine; BA, 6-benzylaminopurine; tZ, *trans*-zeatin; cZ, *cis*-zeatin; Kin, kinetin; TD, thidiazuron; DHZ, dihydrozeatin, Ade, adenine.

**Table 2.** Fold increase of affinity to various cytokinins of mutant receptors compared to ZmHK1.

| Cytokinin | Receptor |  |  |
| --- | --- | --- | --- |
|  | *ZmHK1-AEWL* | *ZmHK1-1603* | *ZmHK1-1595* |
| iP | 8.06 | 4.92 | 2.26 |
| BA | 6.59 | 4.05 | 4.58 |
| tZ | 4.98 | 4.37 | 4.34 |
| cZ | 6.74 | 5.56 | 3.88 |
| Kin | 4.33 | 4.40 | - |
| TD | 0.4 | 0.39 | 0.68 |
| DZ | 5.06 | 3.55 | 8.09 |
| **Assay** | Bacterial spheroplasts |  | Tobacco membrane |

iP, *N*<sup>6</sup>-( $\Delta^2$ -isopentenyl)adenine; BA, 6-benzylaminopurine; tZ, *trans*-zeatin; cZ, *cis*- zeatin; Kin, kinetin; TD, thidiazuron; DHZ, dihydrozeatin, Ade, adenine.

### The *Hsf1* missense mutations localize near the CK binding pocket in ZmHK1

To gain better insight into how each *Hsf1* missense mutation might impact CK binding, we determined the effect these mutations had on the structure of the CHASE domain, which was facilitated by the publication of the crystal structure of the *Arabidopsis thaliana histidine kinase4* (*AHK4*) gene CHASE domain (Hothorn *et al*., 2011). *AHK4* is co-orthologous to *ZmHK1* (69% identical and 83% similar within 245 residues of the CHASE domain) and three other paralogous histidine kinases in the maize genome (Steklov *et al*., 2013). To explore the effects of the *Hsf1* mutations on receptor structure, homology modeling was used first to model the 3D structure of the CHASE domain of ZmHK1 using the structure of AHK4. This was done with and without CK occupying the binding pocket, which did not change the results. Given the high degree of amino acid identity between ZmHK1 and AHK4, the ZmHK1 CHASE domain structure was resolved with high confidence. Next, each mutant receptor was modeled based on the derived ZmHK1 structure. The models were subjected to dynamics simulation with appropriate solvation (see Methods). The results of homology modeling showed that the amino acids mutated in each *Hsf1* allele do not occur within the CK binding pocket (Figure 1F) and thus do not contribute to direct polar contacts with the ligand. Instead, each altered residue is located near a loop domain that forms one face of the binding cavity. An indication of how the mutated residues at these positions might affect ligand binding was provided by the structure model of the ZmHK1-1603 receptor. The residue altered in ZmHK1-1603 is E236, which is predicted to form an ion-pair interaction with R192 located in the loop domain. This polar interaction may help to stabilize the position of the loop domain (Figure 1F). The *Hsf1-1603* mutation converts E236 to K, a negative to positive residue change, which is expected to break the polar interaction with R192 and possibly destabilize the position of the loop due to the nearness of the two positively charged residues. Altering the position of the loop may change the overall conformation of the ligand binding pocket and, thus, account for differences in ligand binding affinities. The missense residues in the other two mutant receptors could potentially alter the conformation of the CK binding pocket via a different mechanism, although our modeling results did not reveal an obvious one.

### Exogenous CK treatment recapitulated the *Hsf1* phenotype

The biochemical and structural analyses suggested the *Hsf1* mutant receptor might be hypersignaling the perception of CK which altered leaf development. To test the idea that increased CK signaling could produce *Hsf1*-like phenotypes, wild type, B73 inbred seeds were transiently treated with the CK 6-benzylaminopurine (6-BAP). The embryo in a mature maize seed possesses about 5 leaf primordia and it is these primordia which experienced the hormone treatment (Kerstetter and Poethig, 1998). Imbibed seeds were treated for 6 days with 10 μM 6-BAP, rinsed and transplanted to soil (see Methods). After growth for 3-weeks, the first 4 seedling leaves were examined for developmental changes (Figures 2A to 2E). Similar to *Hsf1*, 100% of the CK treated B73 seeds produced smaller seedling leaves covered with abundant macrohairs (Figures 2A to 2E). Leaf sheath length, blade length and blade width were reduced by 10% to 20% for leaf 3, similar to leaf size reductions in the *Hsf1* seedlings (Figure 2C). In addition, macrohair density increased on the abaxial sheath, near the auricle, on the adaxial blade, and blade margins in 100% of the CK-treated B73 seedlings (Figures 2D to 2E). This pattern of ectopic macrohair formation was similar to that seen in *Hsf1* seedlings (Bertrand-Garcia and Freeling, 1991a). In addition to alterations in leaf size and pubescence, nearly 20% of the CK treated B73 seeds produced seedlings with prongs on leaf 4 (Figures 2F). This was in contrast to *Hsf1* seedlings where prongs rarely, if ever, developed on leaf 4 (Supplemental Figure 1C). Increasing the concentration of exogenous 6-BAP to 100 μM increased the number of B73 seedlings with prongs on leaf 4 to nearly 90%. (Figure 2F) Thus, transient, exogenous CK treatment recapitulated three prominent aspects of the *Hsf1* phenotype: reduced leaf size, increased macrohair abundance, and formation of prongs on blade margins, confirming these developmental changes can be induced by CK.

**Figure 2.**
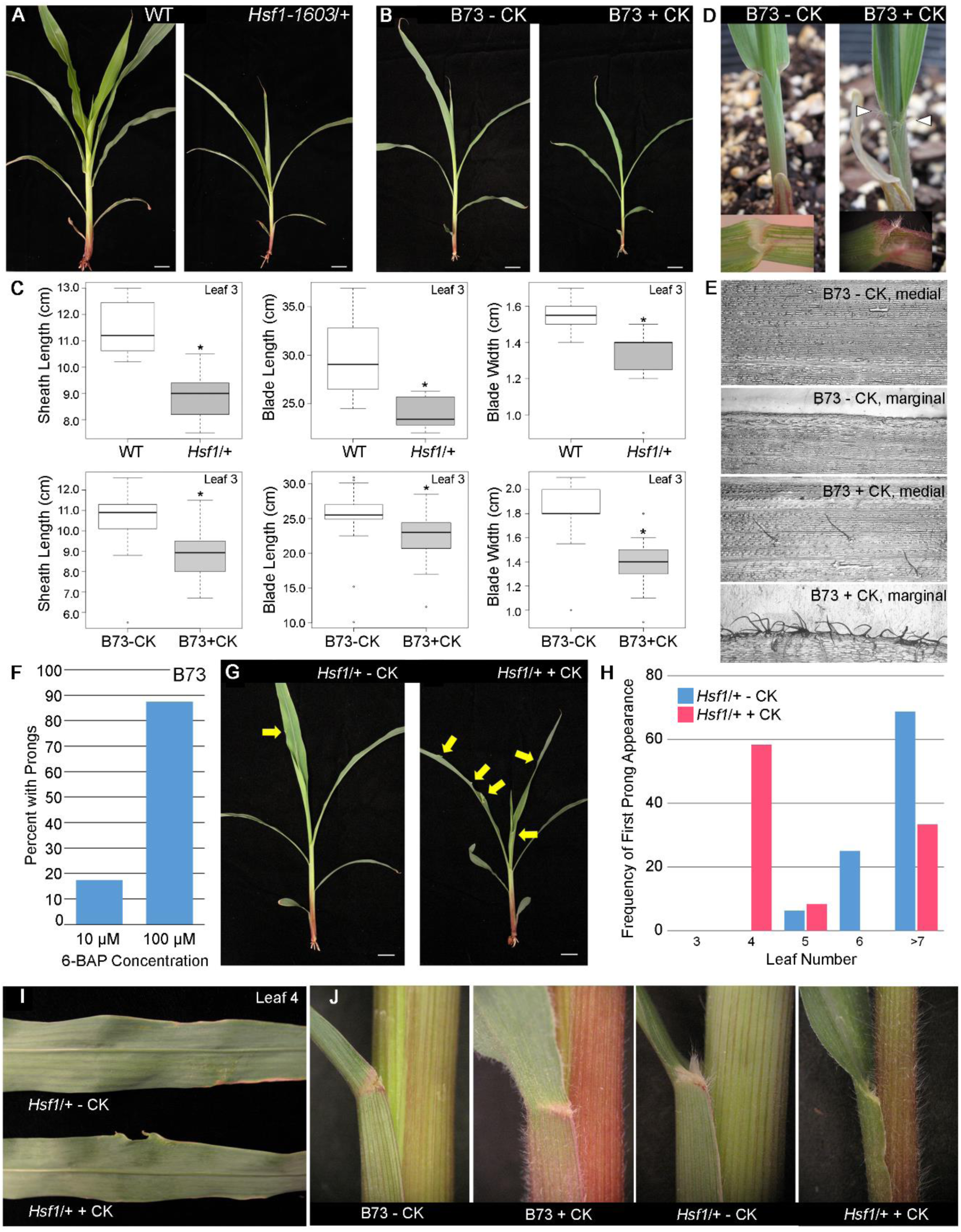
Exogenous CK treatment phenocopies the *Hsf1* leaf development defects and enhances the *Hsf1* mutation. (**A**) Phenotype of 3-week old wild type and heterozygous *Hsf1-*1603/+ seedlings. Bar = 2 cm. (**B**) Phenotypes of 3-week old B73 water (- CK) and 10 µM 6-BAP treated (+ CK) seedlings. Bar = 2 cm. (**C**) Boxplots of leaf sizes comparing wild type (WT) to *Hsf1-1603/+* sib seedlings, and B73 water (- CK) and 10 µM 6-BAP treated (+ CK) seedlings. Horizontal bars represent the maximum, third quantile, median, first quantile, and minimum values respectively, dots outside of the plot are outliers, and the * indicates a *P*-value ≤ 0.0001 calculated from a two-tailed Student’s t-test. (**D**) Macrohair production on the abaxial sheath and auricle (white triangles) of 2-week old B73 water (- CK) and 10 µM 6-BAP treated (+ CK) seedlings. Insets show an adaxial view of the sheath-blade boundary of leaf 1. (**E**) Glue impressions of adaxial leaf 1 blade from 2-week old B73 water (- CK) and 10 µM 6-BAP treated (+ CK) seedlings showing increased macrohair presence in the medial blade and at the margin. (**F**) CK-induced prong formation in B73 seedlings (n ≥ 12 for each treatment). (**G**) Effect of CK treatment on prong formation in 2-week old *Hsf1-1603*/+ seedlings (yellow arrows mark prongs). Bar = 2 cm. (**H**) Frequency and leaf number where the first prong formed in *Hsf1-1603/+* with (red) and without (blue) 10 µM 6-BAP treatment (n ≥ 12 for each treatment). (**I**) Close-up of prongs formed on leaf 4 from CK-treated and control *Hsf1-1603*/+ seedlings (in [G]). (**J**) Macrohair production on 2-week old seedlings due to CK treatment or *Hsf1-1603*/+ mutation or both.

If CK hypersignaling in *Hsf1* was due to increased ligand affinity, then we would expect *Hsf1* to be hypersensitive to CK treatment. To test this idea, we performed six-day treatments on segregating *Hsf1-1603/+* seeds using 0.1 μM CK, a concentration that did not elicit leaf size changes in B73 inbred seed (Supplemental Figure 5A). To distinguish segregating heterozygous *Hsf1* plants from wild type sib plants, PCR genotyping was used to detect a size polymorphism in the *Hsf1-1603* allele (Supplemental Table 3). After CK treatment, seedlings were grown for 3 weeks, after which, leaf phenotypes were measured. While 0.1 μM CK treatment had no effect on wild type sibling leaf size (Supplemental Figure 5A), it did reduce the leaf size of *Hsf1-1603/+* plants 10% to 30% (Supplemental Figure 5B). Thus, *Hsf1-1603/+* seedlings were responsive to a lower concentration of CK that did not elicit a response in wild type sib or B73 inbred seedlings. Treatment with 10 μM 6-BAP was also used to assess effects on prong and macrohair formation in *Hsf1-1603/+* plants. Similar to earlier results (Supplemental Figure 1C), seedlings from control water-treated *Hsf1-1603/+* seeds first formed prongs on leaf 5 (ca. 5%) or leaf 6 (ca. 25%) but never on earlier arising leaves (Figure 2G to 2I). In fact, about 60% of *Hsf1-1603/+* seedlings normally first formed prongs on leaves arising on or after leaf 7 (Figure 2H). By contrast, of the 10 μM 6-BAP treated *Hsf1-1603/+* seeds, nearly 60% produced seedlings where prongs first formed on leaf 4 and only about 30% formed prongs on leaves arising on or after leaf 7 (Figures 2G to 2I). In addition, macrohair abundance appeared increased for CK-treated *Hsf1-1603/+* compared to control *Hsf1-1603/+* or 6-BAP treated wild type sib seedlings but this was not measured (Figure 2J). Thus, CK treatment of *Hsf1* resulted in earlier arising and enhanced mutant phenotypes, indicating the mutation was hypersensitive to the CK hormone, consistent with the biochemical analysis of the mutant receptor.

### CK responsive genes are up-regulated in *Hsf1* leaf primordia

Based on the *Hsf1* mutant plant phenotypes, we presumed that hypersignaling in developing leaf primordia gave rise to the alterations in P-D leaf patterning and other phenotypes. To test this idea, we determined the expression of *ZmHK1* and several CK responsive genes in *Hsf1-1603/+* and wild type sibling plants. Published qPCR and *in silico* expression analyses (https://www.maizegdb.org/gene_center/gene/Zm00001d017977#rnaseq) indicated *ZmHK1* was expressed broadly across several tissues including leaves, roots, stem, and tassel (Yonekura-Sakakibara *et al*., 2004). We reverse transcribed cDNA from three tissues, shoot apices (shoot apical meristem plus 3 youngest leaf primordia), immature leaf, and mature green leaf from two-week old seedlings. Using quantitative PCR (qPCR) we assessed expression in plants heterozygous for the three *Hsf1* alleles compared to their wild type sibs (Figure 3A). We did not detect an increase in *ZmHK1* transcript accumulation in the *Hsf1/+* mutants compared to their wild type controls. Next, we examined expression of CK-responsive genes; two type-A response regulators, *ZmRR3* and *ZmRR6,* and a cytokinin oxidase gene, *ZmCKO4b* (Asakura *et al*., 2003; Giulini *et al*., 2004). We found increased transcript accumulation for all three CK-responsive genes in the *Hsf1/+* mutants, although there was some inconsistencies across genotypes and tissues (Figure 3A).

**Figure 3.**
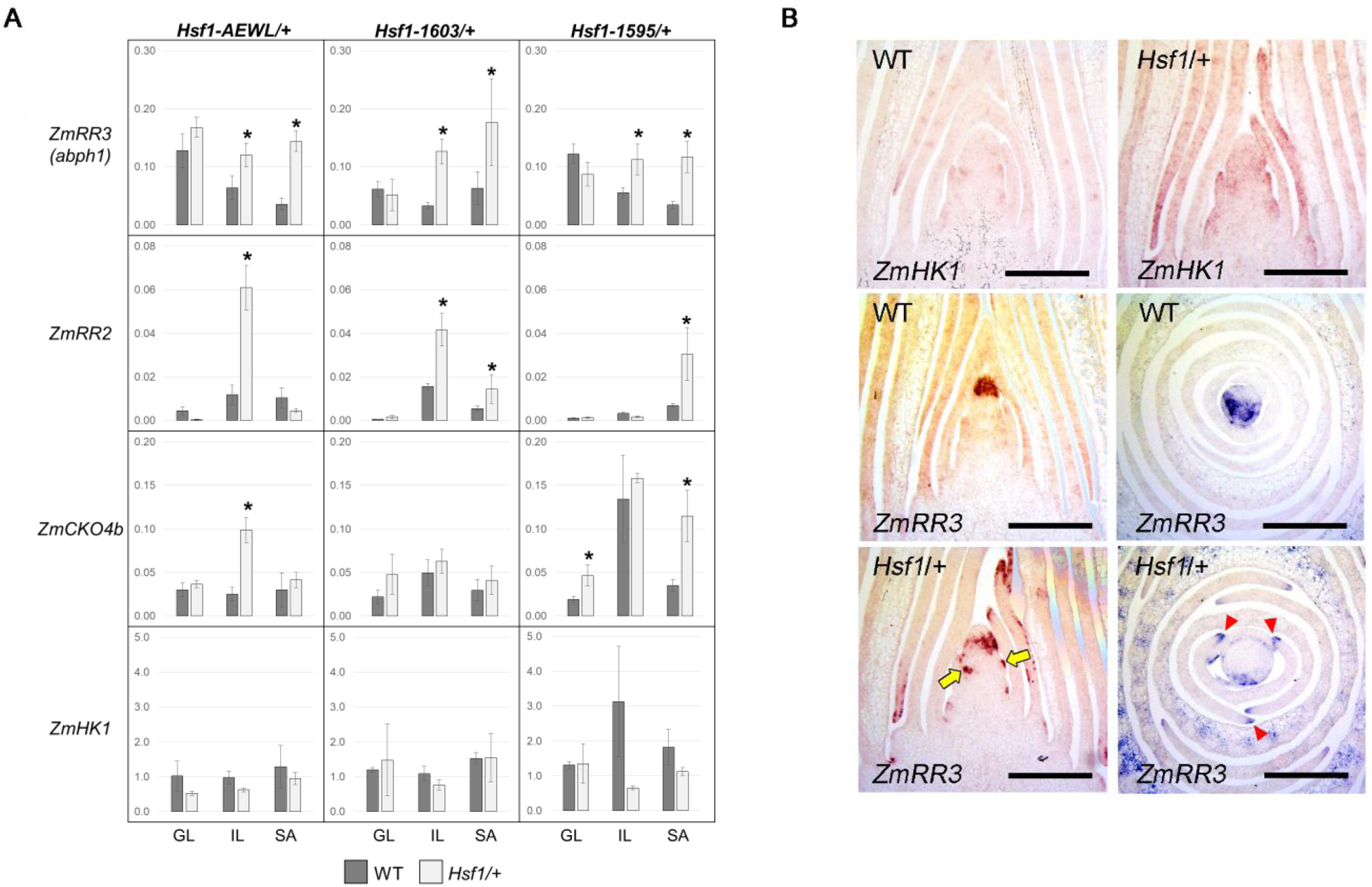
Expression of CK signaling and responsive genes. (**A**) Relative mRNA accumulation of CK genes in different tissues of 2-week old seedlings of the three *Hsf1* alleles and WT sibs measured by qPCR. For each genotype, values are the means (±SE) of three biological replicates consisting of tissue pooled from at least 3 plants. Asterisks indicate significant differences between WT and *Hsf1*/+ sib (Student’s *t* test, P ≤ 0.05). GL – Green leaf, IL – immature leaf, SA –shot apex. (**B**) Pattern of *ZmHK1* and *ZmRR3* transcript accumulation in WT and *Hsf1-1603/+* shoot apex. Longitudinal and transverse sections were hybridized with *ZmHK1* or *ZmRR3* specific antisense probes. The longitudinal section of *ZmRR3* hybridized to WT is not medial and so *ZmRR3* expression appears to be apically localized, but it is not. Initiating leaf primordia (yellow arrows) and leaf primordia margins (red triangles) are marked in the *Hsf1/+* sections probed with *ZmRR3*. Bar = 30 µm.

Using *in situ* hybridization, we assessed transcript localization of *ZmHK1* and *ZmRR3* in wild type and *Hsf1-1603/+* shoot apices (Figure 3B). The *ZmHK1* transcript was found to be distributed broadly within developing leaf primordia and shoot apices in both genotypes (Figure 3B). As was demonstrated previously, *ZmRR3* was expressed in a specific wedge-shaped domain in the apical meristem in both longitudinal and transverse sections of wild type apices but no signal was detected in leaf primordia (Figure 3B) (Giulini *et al*., 2004). However, the spatial expression of *ZmRR3* was expanded in *Hsf1-1603/+* apices. Strong *ZmRR3* expression was visible in its normal meristem domain but signal was also detected in leaf primordia and was particularly evident at the margins (Figure 3B). Given the expanded pattern of *ZmRR3* expression in *Hsf1-1603/+* leaf primordia margins and that *ZmRR3* expression is CK responsive, we interpreted this to indicate increased CK signaling in in the tissue where prongs will form.

### Mutation of *ZmRR3*, a negative regulator of CK signaling, enhances the *Hsf1* phenotype

To test if the increased transcript accumulation of the CK responsive genes was biologically relevant, we made use of a null allele of *ZmRR3*, also known as *aberrant phyllotaxy1* (*abph1*). Plants homozygous for the recessive *abph1* reference allele have an altered phyllotactic pattern and develop leaves paired 180° at each node instead of having the normal alternating pattern (Figures 4A to 4B) but have no P-D patterning defects (Jackson and Hake, 1999). Backcross families were produced which segregated four phenotypes – wild type, heterozygous *Hsf1-1603*, homozygous *abph1*, and heterozygous *Hsf1-1603* plus homozygous *abph1* – in equal frequencies (Figures 4A to 4B). Double mutant plants, heterozygous for *Hsf1* and homozygous for *abph1*, had paired leaf phyllotaxy and a strongly enhanced *Hsf1* phenotype, including very stunted stature, increased shoot branching, very slow growth, extremely short and narrow leaves, and severe leaf patterning defects including abundant prongs and bi- or trifurcation of leaf blades (Figure 4B). The synergistic interaction of *Hsf1* and *abph1* was consistent with *ZmRR3* functioning as a negative regulator of CK signaling and indicated the loss of *abph1* function enhanced the *Hsf1* phenotype.

**Figure 4.**
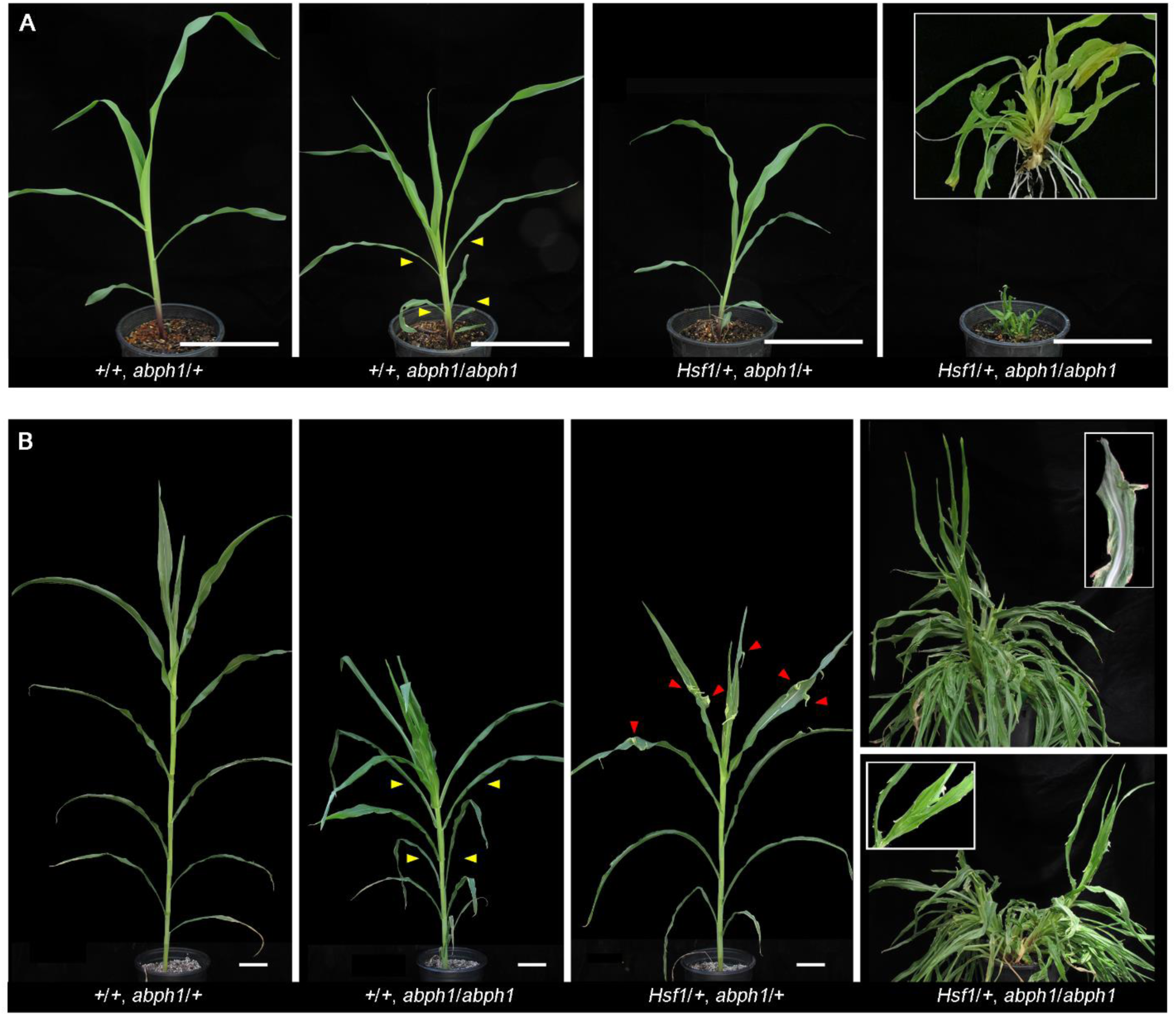
The *Hsf1* phenotype is enhanced by loss of *ZmRR3* function. (**A**) Phenotypes of 30-day old (left to right) WT, *abph1/abph1*, *Hsf1-1603/+*, and *Hsf1-1603/+, abph1/abph1* mutants. This family segregated 9 wild type, 12 *abph1/abph1*, 10 *Hsf1-1603/+*, and 15 double *Hsf1-1603/+*, *abph1/abph1*, which fits a 1:1:1:1 expected ratio. Inset shows a close-up of a double *Hsf1*, *abph1* mutant. Bar = 15 cm. (**B**) Phenotypes of 60-day old plants segregating the same four genotypes in (**A**). Bar = 10 cm. Insets in the double mutant images show close-ups of prongs from that genotype. Yellow and red arrowheads mark paired leaves on the *abph1* mutant and prongs on the *Hsf1/+* mutant, respectively.

## DISCUSSION

### CK influences specific developmental programs in maize leaves

In this study we showed that the *Hsf1* mutation conditions a CK hypersignaling phenotype that has multiple effects on plant growth and development, including specific effects on (i) leaf patterning, (ii) leaf size and (iii) leaf epidermal cell fate (Bertrand-Garcia and Freeling, 1991a). Supporting this idea, we also show exogenous CK treatment of wild type maize seeds produced similar changes in these developmental programs. Prominent among the developmental changes was a specific alteration in P-D leaf patterning where ectopic outgrowths with proximal identity (prongs) formed in the distal blade (Figures 1A to 1C and Supplemental Figure 2A). Although growth along the P-D axis is fundamental to normal leaf development and morphology, its molecular control has not been fully characterized. In eudicots, the activities of several transcription factor genes, such as, *BLADE ON PETIOLE1* (*BOP1*), *LEAFY PETIOLE* (*LEP*), and *JAGGED* (*JAG*), have been linked to the control of P-D leaf development (van der Graaff *et al*., 2000, 2003; Ha *et al*., 2004; Ohno *et al*., 2004; Norberg *et al*., 2005). *BOP* genes have also been shown to influence P-D leaf patterning in monocots like barley and recently, the activity of three, redundant *OsBOP* genes was shown to be required for sheath identity in rice (Tavakol *et al*., 2015; Toriba *et al*., 2019). In several monocots, the misexpression of several class I *knox* genes perturb P-D patterning by potentially altering phytohormone ratios in developing leaf primordia (Reiser *et al*., 2000; Schneeberger *et al*., 1995; Foster *et al*., 1999b; Ramirez *et al*., 2009). Our analysis of *Hsf1*, the second characterized mutation of a maize CK signaling gene, has uncovered a connection between CK and the specification of P-D leaf patterning that is consistent with this hypothesis. How CK drives prong formation is not clear, although the interplay of CK and GA are known to control the degree of leaf complexity in eudicots like Arabidopsis and tomato, through the specification of marginal lobes or leaflets (Jasinski *et al*., 2005; Bar and Ori, 2015). Whether there is any overlap between the mechanism(s) of prong formation in *Hsf1* and leaflet formation in species like tomato will require further analysis. Prong formation itself appears developmentally regulated as prong initiation seems to be coordinated with formation of the ligule suggesting the signals establishing the P-D axis might be transmitted across the entire leaf primordium (Supplemental Figures 1D to 1E). Moreover, prong formation is not random as prongs form only within a certain domain of the blade, with the largest prongs forming more basally (Supplemental Figures 2D to 2F). Intriguingly, this prong-formation region has some overlap with the domain of the leaf blade deleted by mutation of the duplicate *wuschel-related homeobox* (*wox*) genes *narrow sheath1* and *narrow sheath2* (Nardmann *et al*., 2004). This implies that the marginal domain specified by these duplicate *wox* transcription factors may provide a permissive context for prongs to form. This hypothesis can be tested by analysis of prong formation in the triple mutant.

Leaf sheath and blade length, and blade width were reduced in *Hsf1* heterozygotes compared to wild type sib plants at seedling and mature growth stages, consistent with previous reports, and CK treatment recapitulated this phenotype in wild type inbred seedlings (Figures 2A to 2C) (Bertrand-Garcia and Freeling, 1991b). Since CK activity typically promotes cellular proliferation, how CK hypersignaling reduces growth in the shoot is not known, although increased CK signaling is known to reduce root growth (Werner *et al*., 2001, 2003). Typically, reducing CK accumulation or signaling results in smaller leaves and other above ground organs, suggesting increased CK activity might be expected to enhance growth (Werner *et al*., 2001; Nishimura *et al*., 2004). Growth of the maize leaf is organized linearly along its longitudinal axis into distinct zones of cell division, cell expansion and differentiation (Freeling and Lane, 1992). Recent transcriptome, proteome and hormone profiling studies have enumerated multiple regulatory pathways controlling the size of and transitions between the different growth zones, with GA playing a prominent role (Li *et al*., 2010; Nelissen *et al*., 2012; Facette *et al*., 2013). How increased CK signaling impacts these growth zones to determine final leaf size will require further analysis building upon these previous studies.

In addition to a change in P-D patterning and reduction in leaf size, the *Hsf1* mutation and CK treatment of wild type seed promoted increased macrohair formation in the leaf epidermis (Figures 2D to 2E and 2J). Macrohairs are normally found on adult leaves on the abaxial sheath, at high density near the ligule but declining basipetally, on the adaxial blade and along the blade margin. *Hsf1* increased macrohair production not only on the abaxial sheath, adaxial blade, auricle and blade margins of adult leaves but also on juvenile and transitional leaves which are typically glabrous. CK treatment phenocopied the increased pubescence phenotype of *Hsf1* (Figures 2D to 2E). The epidermis of the maize leaf has three types of pubescence – bicellular microhairs, macrohairs and prickle hairs – with macrohairs being the most prominent (Freeling and Lane, 1992). Macrohairs form by differentiation of specialized epidermal cells organized in patterned files beginning in the fifth or sixth leaf (Moose *et al*., 2004). Little is known regarding the signals specifying macrohair formation, although a recessive mutation affecting macrohair initiation, *macrohairless1*, has been reported (Moose *et al*., 2004). By contrast, trichome differentiation in the leaves of eudicots, like Arabidopsis, is known to be controlled by a core network of positive and negative transcriptional regulators (Ishida *et al*., 2008; Grebe, 2012; Pattanaik *et al*., 2014). And trichome initiation on the inflorescence organs in Arabidopsis is jointly stimulated by the activity of CK and GA, and downstream transcription factors (Gan *et al*., 2007; Zhou *et al*., 2013). The increase in macrohair formation mediated by CK treatment or the *Hsf1* mutant suggests CK can reprogram epidermal cell fate in maize leaves as well.

### Missense Mutations in the Maize CK Receptor ZmHK1 underlie the *Hsf1* phenotype

Our data indicate gain-of-function mutations of the maize CK receptor *ZmHK1* underlie the semi-dominant *Hsf1* mutations. CK signaling, which is well described (To and Kieber, 2008; Hwang *et al*., 2012), regulates several developmental and physiological processes, although influences on leaf patterning are not among them. For example, combinations of loss of function mutations of the three Arabidopsis CK receptors demonstrate this gene family has partially overlapping and redundant functions in the control of shoot and root growth, seed size, germination and leaf senescence (Higuchi *et al*., 2004; Nishimura *et al*., 2004; Riefler *et al*., 2006). CK receptors were shown to also possess phosphatase activity by analysis of a specific mutation of *AHK4/CRE1*, the recessive *wooden leg* (*wol*) allele (*CRE1 T278I*) (Mahonen *et al*., 2006). Plants homozygous for the *wol* allele have abnormal root vascular development due to the dose-dependent constitutive phosphatase activity of this allele. A gain-of-function mutation in the CHASE domain of AHK3 (*ore12-1*) revealed this receptor plays a major role in CK-mediated leaf senescence; although how this mutation affected receptor activity was not explored (Kim *et al*., 2006). The study of gain-of-function mutations has revealed additional information on CK receptor function. Novel, dominant, missense mutations in *AHK2* and *AHK3*, the *repressor of cytokinin deficiency* alleles (*rock2* and *rock3*) enhanced CK signaling, increased CK hypersensitivity, and increased transcript accumulation of CK-responsive genes, similar to the *Hsf1* mutations (Figure 3) (Bartrina *et al*., 2017). In contrast, the *rock* mutations had the opposite effect on phenotype compared to *Hsf1*, producing early flowering, enlarged rosette leaves and shoots, and longer roots. The contrasting phenotypic effects might be due to differences in signaling strength between the *rock* and *Hsf1* mutations or reflect differences in the downstream circuitry between the two species.

### Mutations near the CK binding pocket alter ligand affinity and receptor signaling

To clarify how the function of ZmHK1 was altered in the *Hsf1* mutants, we analyzed their activity in heterologous his-kinase signaling and ligand binding assays. Our results indicate two of the *Hsf1* mutant receptors signal independent of added CK in yeast and all three have increased binding affinities for the natural CKs tested (Figure 1E and Table 1). The mutant receptors may be in a “locked on” state, similar to what was hypothesized for the *snf2* mutation or the increased ligand affinities of the *Hsf1* receptors may explain their ability to signal independent of CK action. We favor the second idea and think the increased CK affinity explains the ability of the mutant receptors to signal in heterologous hosts. Many microbes, including *E. coli* and yeast, contain low concentrations of iP as a normal constituent of tRNA which can become free due to tRNA decay (Skoog and Armstrong, 1970; Hall, 1973; Romanov, 1990; Mok and Mok, 2001). The three mutant receptors all have increased affinity for iP (Table 2). This stronger affinity may be due to stronger complex formation, or longer receptor occupancy and, as a consequence, stronger signaling even in the presence of low iP concentration. Thus, the ability of the ZmHK1-AEWL and ZmHK1-1603 receptors to signal in yeast without added CKs may be due to their increased affinity for iP already present at low concentration in yeast cells (Figure 1E). In fact, it has been shown that expressing other HK receptors in the *sln1* deletion yeast strain TM182 permits this strain to grow on glucose without added CKs, albeit at a much slower rate than with CKs present, and recombinant HKs synthesized in *E. coli* cannot be crystalized without iP complexed in the binding pocket (Higuchi *et al*., 2009; Hothorn *et al*., 2011). Since all three mutant receptors have increased ligand affinities (Table 1), have nearly identical mutant plant phenotypes in several different genetic backgrounds (Figure 1 and Supplemental Table 1), and show similar misexpression patterns of CK responsive genes (Figures 3A) we conclude all three *Hsf1* mutant receptors function similarly *in planta*.

Our structural analysis localized each residue mutated in *Hsf1* to the ligand-binding Per-Arnt-Sim-like (PAS) subdomain of the CHASE domain in ZmHK1 (Figure 1F) (Steklov *et al*., 2013; Hothorn *et al*., 2011). Notably, none are within the CK binding pocket or predicted to make contact with the ligand. Rather all are located near a loop domain comprising one face of the pocket suggesting interactions with this loop may affect the binding pocket resulting in increased ligand affinity. Interestingly, amino acid substitutions that rendered *AHK4* constitutively active in a heterologous *E*. *coli* his-kinase assay were located downstream of the CHASE domain in the second transmembrane domain and near the kinase domain (Miwa *et al*., 2007). In addition, none of the *rock* mutations are located in the ligand-binding PAS domain (Bartrina *et al*., 2011). Rather two are in the N-terminal α–helices and one is in the C-terminal transmembrane domain. Therefore, further structure-function studies will be needed to define which residues are crucial for activity and to resolve the precise mechanism(s) by which individual missense mutations alter ligand binding and receptor signaling.

### *Hsf1* affects downstream components of CK signaling

More ZmHK1 signaling in developing *Hsf1* leaf primordia resulted in increased transcript accumulation of several early CK response genes in all three *Hsf1* mutant alleles (Figure 3A). Although not all CK reporters responded the same within an allele or tissue, overall our data are consistent with *Hsf1* upregulating CK responsive genes. The most consistent effect was upregulation of *ZmRR3* where its normally meristem-confined expression was expanded in *Hsf1-1603* to include expression near newly arising leaf primordia and in primordia margins (Figure 3B). Notably, the increased CK signaling reported by *ZmRR3* marks the margins of early stage leaf primordia (Figure 3B) which is where prongs will form later in development (Supplemental Figures 1F to 1G). Although we found ectopic *ZmRR3* signal along the entire margin, outgrowths do not emanate from the entire blade margin but, rather, occur sporadically, with outgrowths interspersed with regions of normal blade margin (Figures 1B to 1C and Supplemental Figure 2A). This observation suggests even though CK hypersignaling can promote proximalization of blade margin cells, not all cells at the margin are competent to respond to this signal. Double mutants heterozygous for *Hsf1-1603* and homozygous for *abph1*, a null allele of *ZmRR3*, show a synergistic interaction (Figures 4A to 4B). Several type-A RRs function to negatively regulate CK signal transduction, as well as, regulate circadian rhythms, phytochrome function and meristem size (To *et al*., 2004). The increased severity of growth defects in *Hsf1* heterozygotes which lack *abph1* activity suggests upregulation of *ZmRR3* (*abph1*) partially ameliorates CK hypersignaling. This also suggests that *ZmRR3* normally functions to attenuate CK signal transduction in maize shoot apices, in addition to specifying leaf phyllotaxy.

The identification of the CK receptor *ZmHK1* as the gene underlying the leaf patterning *Hsf1* mutation adds to our understanding of the role CK can play in basic developmental programs. Future studies to determine the molecular determinants functioning downstream of CK signaling that promote prong formation should illuminate mechanisms important for developmental reprogramming and cell fate acquisition.

## METHODS

### Plant Material, Genetics, Phenotypic Measurements, and Analysis

The *Hsf1-1595*, *Hsf1-1603* and *Hsf1-2559* mutants arose via EMS mutagenesis of the inbred Mo17 and seed was obtained from the Maize Genetic Cooperation Stock Center (http://maizecoop.cropsci.uiuc.edu/). *Hsf1-AEWL* arose via EMS mutagenesis of the inbred A619 and *Hsf1-7322* via EMS mutagenesis of the inbred A632 in independent screens. Homozygous *Hsf1* mutants of all five alleles were identified for sequence analysis from progeny of self-pollinated heterozygous B73 introgressed plants by phenotype and also by PCR screening of linked sequence polymorphisms unique to the progenitor inbred lines and the backcross inbred B73. Since *Hsf1-1595* and *Hsf1-1603* were the same transition mutations as *Hsf1-7322* and *Hsf1-2559*, respectively, further analysis was only performed on three mutants: *Hsf1-1595*, *Hsf1-1603* and *Hsf1-AEWL*. All phenotypic, molecular and epistatic analyses were performed on the three alleles that had been backcrossed a minimum of six times to the inbred B73. The *Hsf1* phenotype of the three alleles was fully penetrant as a heterozygote in all backcross generations. Progeny from self- or sib-pollinated *Hsf1* heterozygotes of the three alleles segregated 25% severely stunted, very slow growing, multi-shoot plants that only survived when grown in the greenhouse but were sterile. The *abph1* mutant seed was backcrossed a minimum of three times to the inbred B73 before making the double mutant family segregating with *Hsf1-1603*. *Hsf1-1603* heterozygotes were crossed by *abph1* homozygotes and double heterozygous progeny plants were backcrossed by *abph1* homozygotes creating double mutants families segregating 25% +/+, *abph1*/+ (WT); 25% +/+, *abph1*/*abph1* (single *abph1* mutant); 25% *Hsf1*/+, +/*abph1* (single *Hsf1* mutant); and 25% *Hsf1*/+, *abph1*/*abph1* (double *Hsf1 abph1* mutant). Allele specific PCR genotyping was done to confirm phenotypes of *Hsf1* heterozygotes and *abph1* heterozygotes and homozygotes (Supplemental Table 3).

Measurement of adult plant traits of the three *Hsf1* mutant alleles was performed on field grown families segregating 50% wild type: 50% *Hsf1* heterozygotes. Measurements were taken on 7-11 plants of each genotype in 1-row plots with two biological replicates. For analysis of prong position, prong size and percent prong margin, the third leaf above the ear of adult *Hsf1* heterozygous plants was collected from 1-row plots of field grown plants in three replicates in summer 2013. Approximately, 6 to 10 leaves were collected per plot for each allele. For each leaf, measurements were made for (1) total blade length, (2) prong position by measuring the distance from the base of the blade to the mid-point of each prong on each blade margin, and (3) prong size by measuring from the basal to the distal position along the margin where proximal tissue emerged from the blade for each prong (Figure 1B). Percent prong margin was defined as the proportion of leaf blade margin that is occupied by tissue having proximal (sheath, auricle and/or ligule) identity and was calculated by summing the size of all prongs from both sides of the leaf blade divided by twice the length of the leaf blade.

Analysis of prong position, prong size and the relationship between prong position and size was estimated with kernel smoothing methods (Silverman, 1986; Wand and Jones, 1995). For all cases a Gaussian kernel was used and the data reflection method was applied for boundary correction since both position and size are positive variables. The bandwidth were selected using least squares cross validation (Bowman, 1984). All computations were performed using R software, kernel density estimation was performed using the ks package (Duong, 2007) and figures were created with the ggplot2 package (Wickham, 2009).

### Map-based cloning of *Hsf1*

*Hsf1-1595* was introgressed into B73 and crossed to PRE84 to generate a BC1 mapping population. Genetic mapping with 96 BC1 individuals defined *Hsf1* between two SNP markers on chr5: PHA12918-F (204590502 bp, B73 RefGen_v2) and PHA5244-F (206614542 bp, B73 RefGen_v2). The two flanking markers were used to screen a BC1 population of 1500 individuals from B73_*Hsf1* x A632 and 1600 individual from B73_*Hsf1* x PRE84. 224 recombinants were identified, and these individuals were used for further fine mapping. Additional markers derived from the *Hsf1* interval were developed and used to fine map the *Hsf1* mutation with the recombinants, as described in Jiang *et al*., 2012 (Jiang *et al*., 2012). The gene underlying the *Hsf1* mutation was finally delimited to a 21 kb interval, between Indel marker 410984 (205538463 bp, B73 RefGen_v2, with one recombinant between this marker and *Hsf1*) and SNP marker 391087 (205559234 bp, B73 RefGen_v2, with three recombinants between this marker and *Hsf1*). There is only one annotated gene model (B73 RefGen_v3 GRMZM2G151223, B73 RefGen_v4 Zm00001d017977) in this interval that was also annotated in NCBI as LOC541634 histidine kinase1, a putative cytokinin receptor.

### Heterologous histidine kinase assays

Signaling of the wild type and *Hsf1* mutant ZmHK1 receptors in yeast was performed as described previously (Inoue *et al*., 2001). The exact point mutations for each of the three *Hsf1* missense mutations were engineered into the cDNA of *ZmHK1* in the plasmid P415-CYC1-ZmHK1 plasmid with the QuikChange II Site-Directed Mutagenesis kit (Agilent Technologies) using the manufacturer’s specifications.

### Cytokinin binding affinity determination

Cytokinin binding assays were performed with recombinant maize cytokinin receptors expressed in *E. coli* cells. Spheroplasts were prepared from cell lines expressing the wild type ZmHK1, and mutant ZmHK1-AEWL and ZmHK1-1603 receptors. Competitive cytokinin binding assays were performed as previously described (Lomin *et al*., 2011). Transient expression of receptors for the homologous binding assay was done by transformation of tobacco *Nicotiana benthamiana* as previously described (Sparkes *et al*., 2006). Agrobacteria *A. tumefaciens* carrying cytokinin receptor genes fused to GFP were grown in parallel with a helper agrobacterial strain p19 (Voinnet *et al*., 2003). Five to six week old tobacco plants were infiltrated with the mixture of two agrobacterial strains and the expression level of receptor genes was checked after 4 days using a confocal microscope. For those cases with sufficient expression, leaves were processed further for plant membrane isolation. For plant membrane isolation, all manipulations were done at 4 °C. Tobacco leaves were homogenized in buffer containing 300 mM sucrose, 100 mM Tris-HCl (pH 8.0), 10 mM Na2-EDTA, 0.6% polyvinylpyrrolidone K30, 5 mM K2S2O5, 5 mM DTT and 1 mM PMSF. The homogenate was filtered through Miracloth (Calbiochem), and the filtrate was first centrifuged for 10 min at 10000 g, and then for 30 min at 100000 g. The microsome pellet was resuspended in PBS (pH 7.4), frozen and stored at −70 °C before using.

### ZmHK1 structure modeling

The amino acid sequence of the ZmHK1 CHASE domain (86-270) was obtained from the protein sequence database of NCBI (accession id: NP_001104859). It shares 69% sequence identity with the *Arabidopsis HK4* sensor domain. The homology model for ZmHK1 was generated using Swiss model server (http://swissmodel.expasy.org) with the published crystal structure of AHK4 (pdb code: 3T4J) as a template. Subsequently the model was solvated and subjected to energy minimization using the steepest descent followed by conjugate gradient algorithm to remove clashes. The stereochemical quality of the ZmHK1 model was assessed using the PROCHECK program. None of the residues were in the disallowed regions of the Ramachandran map.

### Exogenous CK treatment

Exogenous CK treatments were performed with 6-benzylaminopurine (6-BAP) (Sigma Aldrich) dissolved in 10 drops 1N NaOH and brought to 1mM concentration with distilled water. All water control treatments were done using a similar stock of 10 drops 1N NaOH and diluted in parallel to the CK stock. Further dilutions to the desired CK concentration were done with distilled water. Maize kernels were surface sterilized with two 5 minute washes of 80% ethanol followed by two 15 minute washes of 50% bleach and rinsed five times in sterile distilled water. Kernels were imbibed overnight with sterile distilled water prior to the start of the hormone treatment. For hormone treatments, 20 imbibed kernels per replicate were placed embryo-side down on two paper towels in a petri dish, covered with two more layers of paper towel and filled with 15 mL of CK treatment or the water control solution. Petri dishes were sealed with parafilm and placed in a lab drawer in the dark at room temperature for 6 days. After treatment, germinating kernels were rinsed with sterile, distilled water and planted in 4 cm square pots in soilless potting medium (Metro-Mix 900, SunGro Horticulture) and grown in the greenhouse (day: 16 hr./28°C, night: 8 hr./21°C) with supplemental lighting (high pressure sodium and metal halide lights) and standard light intensity (230 µE m^-2^ s^-1^ at height of 3.5 feet). Growth was monitored and leaf measurements were taken after the fourth leaf collar (auricle and ligule) had fully emerged from the whorl after 3 to 4 weeks. For measurements, individual leaves were removed from the plant and each component measured. Leaf sheath length was defined as the site of insertion of the leaf base to the culm (stem) to the farthest point of sheath adjoining the ligule. Leaf blade length was defined as the most proximal point of blade adjoining the ligule to the distal blade tip. Leaf blade width was measured margin to margin at half of the leaf blade length. All leaf measurements were analyzed using JMP PRO 12 software using a student’s t-test to determine significance with two comparisons, and Tukey’s HSD test to determine significance with more than two comparisons. To examine macrohair abundance, epidermal impressions were made using Krazy Glue Maximum Bond® cyanoacrylate glue applied to a Fisherbrand Superfrost Plus® microscope slide. The adaxial blade of leaf one was pressed firmly into the glue for about 30 seconds, followed by immediate removal of the leaf. Slides were imaged on an Olympus BX60 light microscope.

### Expression analysis

#### *In situ* hybridization

For *in situ* hybridization, we slightly modified an online protocol from Jeff Long. For complete details refer to http://pbio.salk.edu/pbiol/in_situ_protocol.html. *In situ* probes were made using T7/SP6 promoter based *in vitro* transcription in the cloning vector pGEMT (Promega). FAA (Formaldehyde Acetic Acid) fixed and paraffin embedded maize shoot apices were sectioned at 10μm thickness and laid on Probe-On-Plus slides (Fisher) and placed on a warmer at 42°C. After overnight incubation, the slides were deparrafinized using Histo- Clear (National Diagnostics), treated with proteinase K and dehydrated. Probes were applied on the slides and pairs of slides were sandwiched carefully and incubated at 55°C overnight. The following day the slides were rinsed and washed. Diluted (1:1250) anti-DIG-antibody (Roche) was applied to the slides and incubated for 2 hours. After thoroughly washing the slides, sandwiched slides were placed in NBT-BCIP (Roche) solution (200 μl in 10ml buffer C; 100mM Tris pH9.5/100mM NaCl/50mM MgCl_2_) in dark for 2 to 3 days for color development. Color development reaction was stopped using 1x Tris EDTA. The slides were mounted using Immu-Mount (Thermo Scientific) and observed and imaged under a bright field microscope.

#### RT qPCR

Seedling tissue was collected from two-week old, stage V3 to V4 *Hsf1/*+ and wild type sib seedlings for each allele and included (1) ca. 2 cm of mature green leaf blade from the distal half of leaf #4, (2) ca. a 2 cm cylinder of immature leaf tissue, cut ca. 1 cm above the insertion point of leaf #5 after removing leaf #4, and (3) the remaining 1 cm cylinder of tissue above the insertion point of leaf #5, consisting of the SAM, young leaf primordia and the apical part of the stem. Tissue was bulked from three different plants for each biological replicate and three replicates were collected. Total RNA was extracted from these tissues using Trizol reagent, adhering to the manufacturer’s protocol (http://tools.lifetechnologies.com/content/sfs/manuals/trizol_reagent.pdf). cDNA was synthesized from total RNA using the SuperScript® III First-Strand (Invitrogen) synthesis system for reverse transcriptase PCR (RT-PCR) and oligo-d(T) primers. Quantitative real-time PCR was performed on the cDNA using an LC480 (Roche) and the SYBR green assay. The primers were designed near the 3’ end of the gene with an amplicon size of between 120 bp to 250 bp. Folylpolyglutamate synthase (FPGS) was used as an endogenous control as it was shown to have very stable expression across a variety of maize tissues and range of experimental conditions (Manoli *et al*., 2012). Two technical replicates were included for each gene. Comparative ΔΔCt method was used to calculate fold change compared to the endogenous control. ΔCt of mutant (*Hsf1*) and ΔCt of wild type (WT) was expressed as the difference in Ct value between target gene and the endogenous control. ΔΔCt was then calculated as the difference of ΔCt (*Hsf1*) and ΔCt (WT). Finally, fold change in target gene expression between *Hsf1* and WT was determined as 2 ^−ΔΔCt^.

## Acknowledgements

We thank Dave Jackson (Cold Spring Harbor Labs) for the *abph1* mutant seed, Erica Unger-Wallace (Iowa State University) for technical assistance and Erik Vollbrecht (Iowa State University) for support during the early phase of this research. D.M.K., S.N.L. and G.A.R. were partly supported by the Molecular and Cell Biology Program of the Presidium of RAS. This work was supported by the National Science Foundation under Grant Number 1022452.

## Author contributions

M.G.M., S.C., H.S., G.A.R., B.L. and N.B. designed research; L.M.T., S.C., J.C., A.D.V.E., I.A., A.P., D.M.K., S.N.L., S.T., and N. M. performed research; M.G.M., H.S., G.A.R., B.L. and N.B. analyzed data; and M.G.M. wrote the paper with input from the other authors.

## Supplemental Data

**Supplemental Figure 1.** Hsf1 phenotypes.

**Supplemental Figure 2.** Prong formation is patterned in Hsf1 leaves.

**Supplemental Figure 3.** ZmHK1 activity in heterologous yeast his-kinase assay.

**Supplemental Figure 4.** Comparison of ligand binding affinity constants of wild type and mutant ZmHK1 receptors.

**Supplemental Figure 5.** Effects of CK treatment on leaf growth.

**Supplemental Table 1.** Mature plant phenotypes of the three Hsf1 alleles.

**Supplemental Table 2.** Frequency of prongs for the three Hsf1 alleles by leaf position in the upper shoot..

**Supplemental Table 3:** Primers used for positional cloning or genotyping

**Supplemental Table 4.** Primers used for expression analysis

