## Supplemental Data for "The maize *Hairy Sheath Frayed1* (*Hsf1*) mutant alters leaf patterning through increased cytokinin signaling"

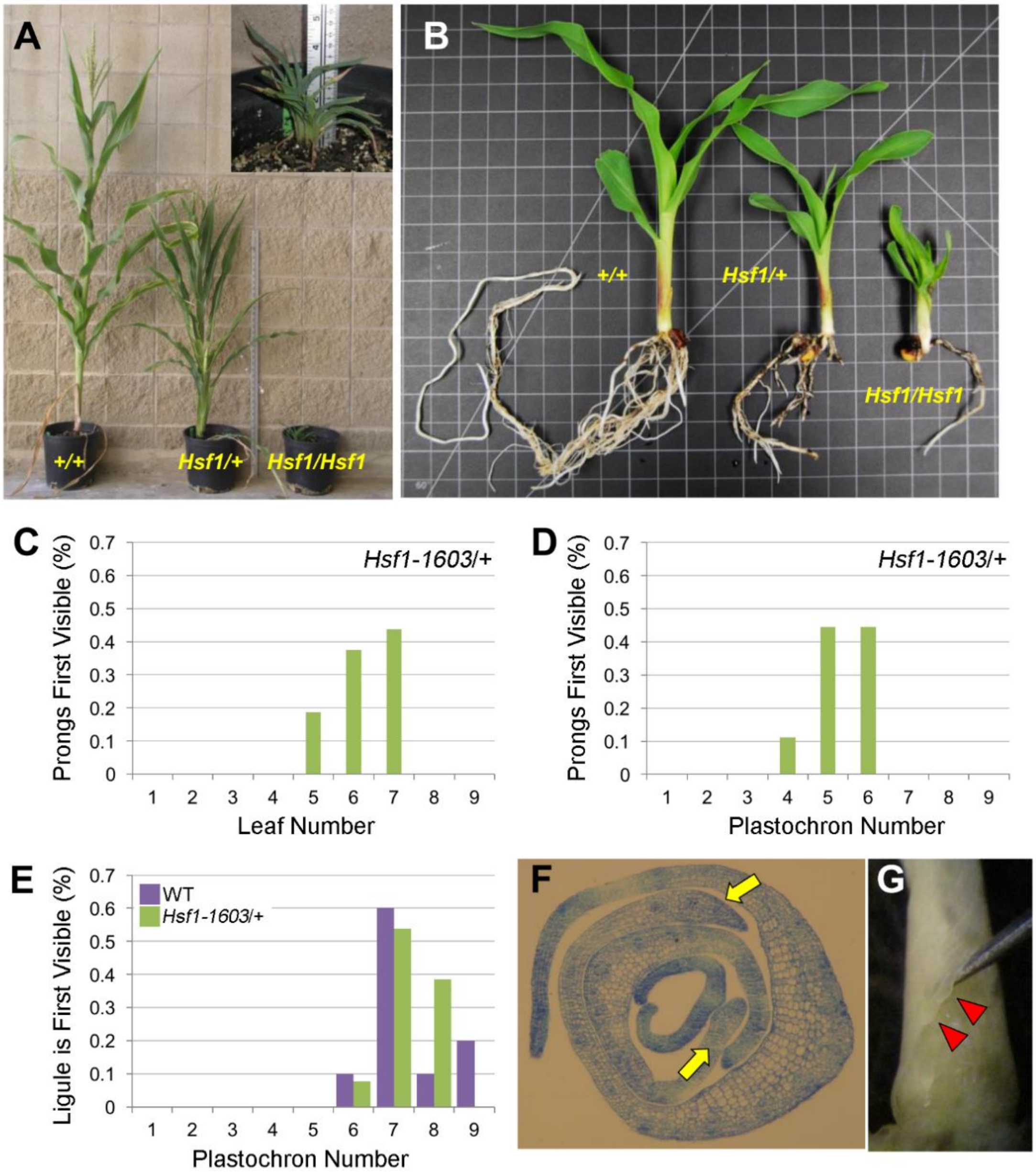


**Supplemental Figure 1. *Hsf1* phenotypes. (A)** Comparison of wild type (+/+), *Hsf1-1603/+* and *Hsf1-1603/Hsf1-1603* sib plants at flowering of wild type. Inset shows close-up of the *Hsf1-1603* homozygote at a comparable growth stage to the other two genotypes. **(B)** Two-week old seedlings of the same genotypes shown in **[A]** showing typical shoot and root phenotypes. **(C - E)** Characterization of prong formation in *Hsf1-1603/+*. Frequency of the leaf number in which prongs first form **(C)**, the plastochron stage where prongs are first visible at the blade margin **(D)**, and the plastochron stage the developing ligule is first visible in 3-week old *Hsf1-1603/+* and wild type sib plants **(E)**. **(F - G)** Examples of developing prongs (yellow arrows), note difference in thickness of margins from same leaf, in a transverse section through P5 and P6 leaf primordia **(F)** and in a hand-dissected whole P6 leaf (red triangles, note sinuate margin) **(G)** both from 25-day old seedlings. All analyses in **(C)** to **(E)** were done on heterozygous *Hsf1-1603* plants in the B73 inbred (≥ 6 backcrosses) from a sample of n = 16 **(C)**, n = 10 **(D)**, or n ≥

10 for both mutant and wild type sibs **(E)**. The stick is 1 meter in **(A)** and the squares are 0.5 inch in **(B)**.

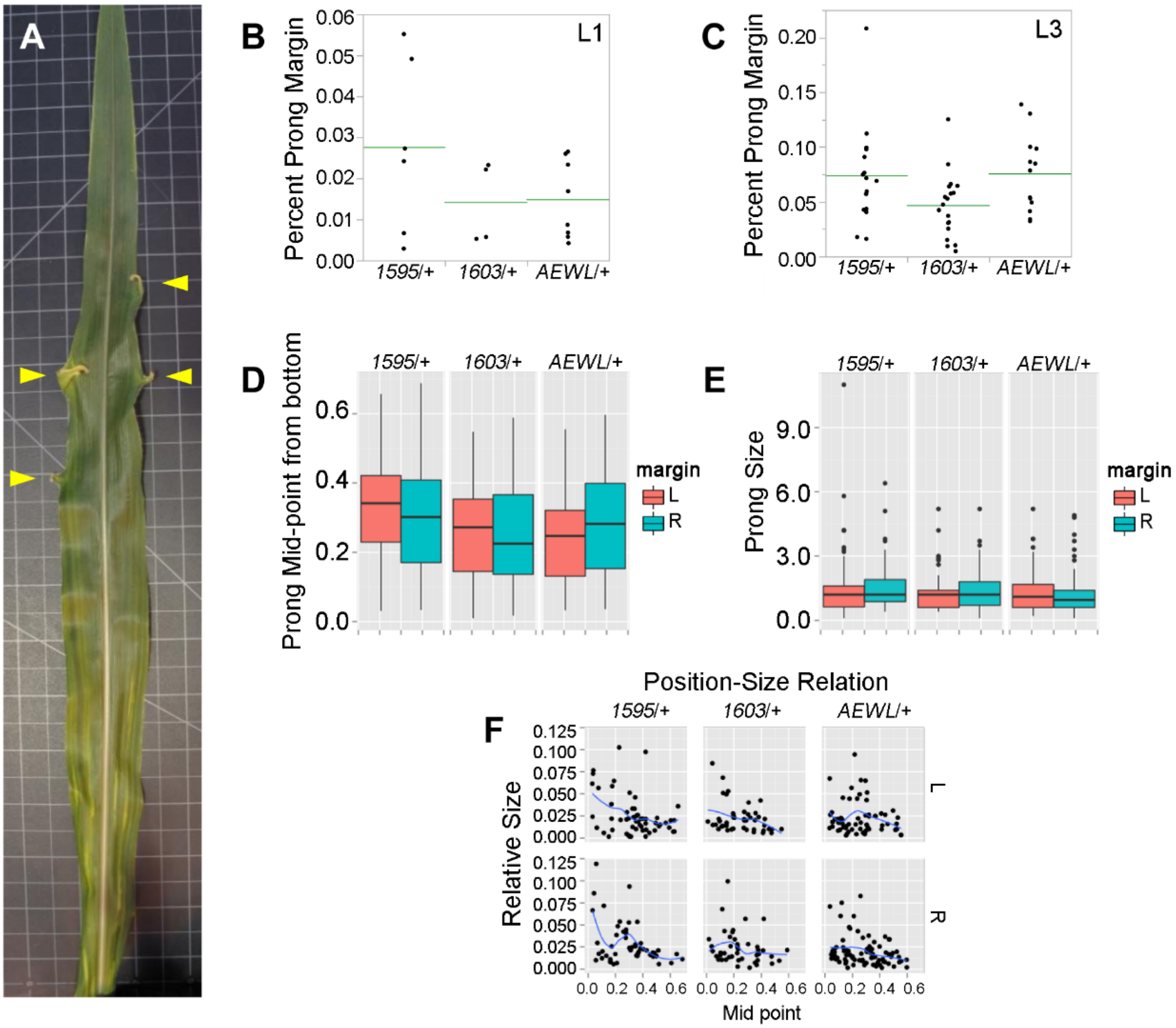


**Supplemental Figure 2.  Prong formation is patterned in *Hsf1* leaves. (A)** Adaxial view of leaf # 6 from *Hsf1-1603*/+ showing the arrangement of prongs (yellow triangles) on both margins.  Squares are 0.5 in.  **(B - C)**  Prong size increases on upper leaves compared to leaves lower on the shoot for the three *Hsf1* alleles.  Scatterplot of the percentage of blade margin occupied by prong tissue (Percent Prong Margin, PPM) on the first leaf (L1) above the top ear **(B)** and on the third leaf (L3) above the top ear **(C)**.  The green lines are means and the points are PPM from different plants.  **(D - E)**  Size and position of prongs along the adaxial-facing blade.  Boxplot analysis of the position along the blade where prongs form as a percent of total blade length **(D)** and prong size in cm **(E)** for the three *Hsf1* alleles.  For boxplots, the box defines the interquartile range, the horizontal line is the median, and the whiskers define the maximum and minimum values.  **(F)** Scatter plot showing the relationship between prong size and prong position on the left (L) and right (R) blade margin for the three *Hsf1* alleles.  Larger prongs form more often in the lower 20% of the leaf blade on both margins.

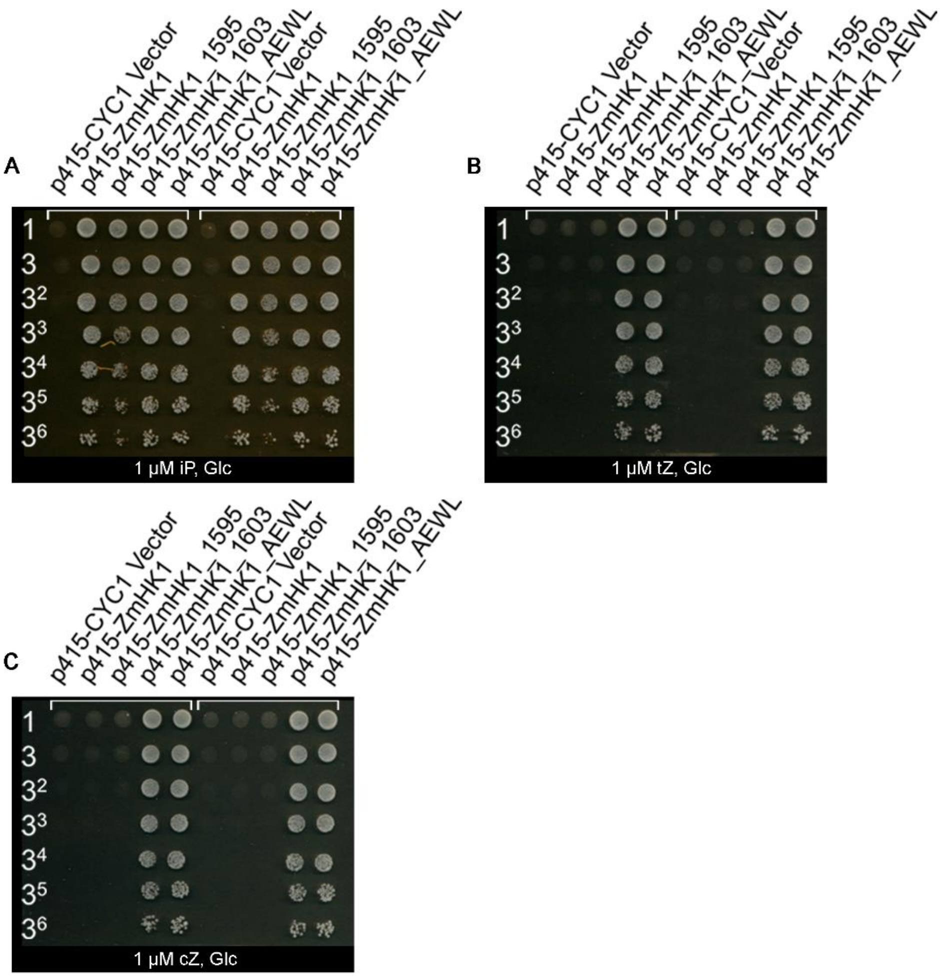


**Supplemental Figure 3.  ZmHK1 activity in heterologous yeast his-kinase assay.  (A - C)** Growth of *S. cerevisiae sln* Δ mutant on glucose media with wild type and mutant ZmHK1 receptors supplemented with low concentration of CKs.  Growth on glucose with 1 µM iP **(A)**, 1 µM tZ **(B)** or 1 µM cZ **(C)**. Dilutions of yeast cultures (O.D.$_{600}$ = 1.0) for each yeast strain are noted on the left of each image.  .  DMSO, dimethyl sulfoxide; iP, $N^6$-($\Delta^2$-isopentenyl)adenine; BAP, 6-benzylaminopurine; tZ, *trans*-zeatin; Kin, kinetin; cZ, *cis*-zeatin.

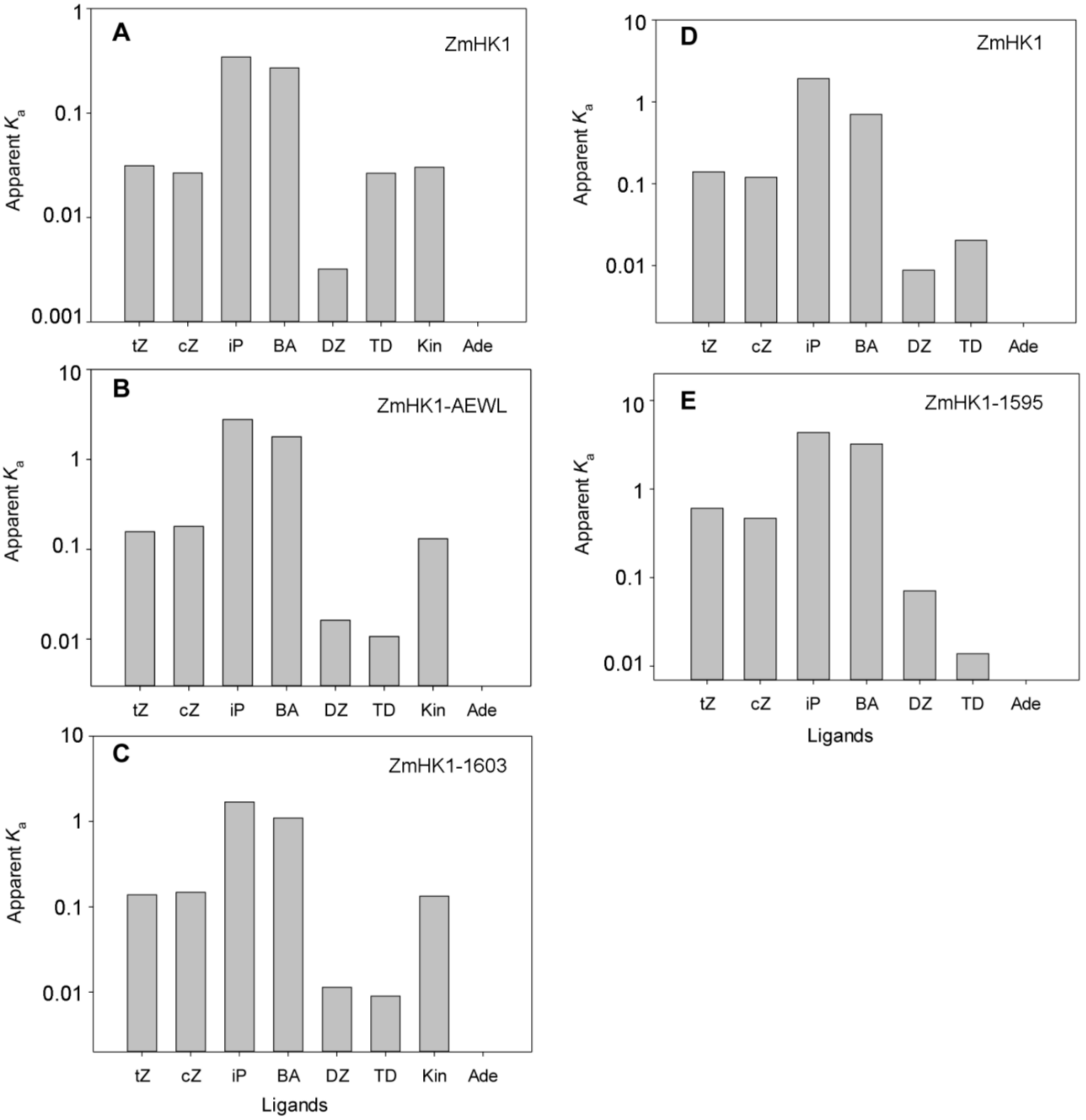


**Supplemental Figure 4. Comparison of ligand binding affinity constants of wild type and mutant ZmHK1 receptors**. **(A - C)** Affinity constants of ZmHK1 **(A)** and mutants ZmHK1-AEWL **(B)** and ZmHK1-1603 **(C)** derived from heterologous *E. coli* assay system using bacterial spheroplasts. **(D - E)** Affinity constants of ZmHK1 **(D)** and mutant ZmHK1-1595 **(E)** derived from homologous tobacco assay using transient expression in *N. Benthamiana* membranes. For all graphs, values of apparent $K_a = K_i^{-1}$ ($nM^{-1}$) are shown.

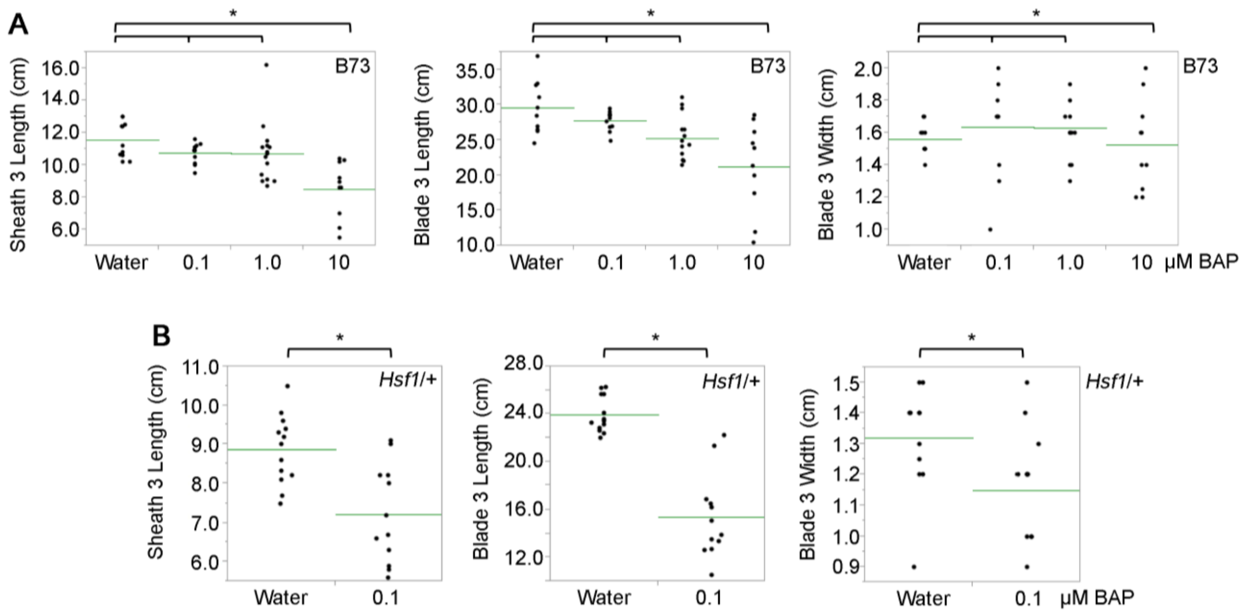


**Supplemental Figure 5.  Effects of CK treatment on leaf growth.  (A)** Growth response of seedling leaf #3 of B73 inbred seed germinated in the presence of various CK concentrations for 6 days.  Note concentrations less than 10 µM do not affect leaf growth in this assay.  **(B)** Growth response of seedling leaf #3 of *Hsf1-1603/+* seed germinated in 0.1 µM CK for 6 days.  Note this concentration of CK has no effect on B73 leaf growth.  In all graphs, each dot represents one leaf measured, the green bar represents the mean and a bracket with a * indicates the compared means are significantly different at a *P* value ≤ 0.05 calculated from a two-tailed Student's *t*-test.

**Supplemental Table 1.** Mature plant phenotypes of the three *Hsf1* alleles.

| Genotype | LFNO | PLHT (cm) | EARSH# | EARLF# | EARLFL (cm) | EARLFW (cm) | TASLEN (cm) |
|---|---|---|---|---|---|---|---|
| **+/+ (*Hsf1-AEWL*)** | 19.5 | 171.1 | 2.5 | 14.4 | 78.2 | 8.2 | 44.6 |
| ***Hsf1-AEWL*/+** | 20.6 | 142.1 | 1.1 | 15.0 | 70.2 | 5.6 | 29.3 |
| **+/+ (*Hsf1-1603*)** | 19.3 | 156.5 | 2.2 | 14.1 | 76.4 | 8.1 | 39.9 |
| ***Hsf1-1603*/+** | 21.0 | 123.3 | 1.1 | 15.6 | 62.9 | 5.4 | 30.3 |
| **+/+ (*Hsf1-1595*)** | 19.9 | 160.2 | 2.1 | 14.8 | 72.8 | 8.2 | 30.1 |
| ***Hsf1-1595*/+** | 21.5 | 128.9 | 1.1 | 16.3 | 63.7 | 6.3 | 28.7 |

All measures are means and all the phenotypic differences except those with gray shading are significant at $P \leq 0.01$ calculated from a two-tailed Student's *t*-test. LFNO – total leaf number, PLHT – plant height, EARSH# - number of visible ear shoots, EARLF# - the number of the leaf subtending the uppermost ear shoot, EARLFL – the length of the leaf subtending the uppermost ear shoot, EARLFW – the width of the leaf subtending the uppermost ear shoot, and TASLEN – length of the central rachis of the tassel. See Methods for experimental details.

**Supplemental Table 2.** Frequency of prongs for the three *Hsf1* alleles by leaf position in the upper shoot.

| | Percentage of leaves with prongs | | |
|---|---|---|---|
| **Genotype** | **1st leaf above ear** | **2nd leaf above ear** | **3rd leaf above ear** |
| *Hsf1-AEWL/+* | 40.0 | 73.7 | 84.2 |
| *Hsf1-1603/+* | 19.0 | 47.6 | 90.5 |
| *Hsf1-1595/+* | 31.6 | 73.7 | 84.2 |

Number of leaves counted was between 19 – 21 for each leaf position and each allele.

**Supplemental Table 3:** Primers used for positional cloning or genotyping

| Primer | Sequence (5' > 3') | Purpose | Notes |
|---|---|---|---|
| PHA12918-F | CCAGTTGCTTCACTTTGTATTA | SNP markers for initial screening | Can be used as CAPS marker using enzyme *Hph*I |
| PHA12918-R | TCATCAAATCAGAAGGGAG | | |
| PHA5244-F | CCATGCAAGTGGTTGGTGC | SNP markers for initial screening | Can be used as CAPS marker using enzyme *Sau*961 |
| PHA5244-R | TGGTTCAAGCGTTCCACAGT | | |
| 410984-F | GAGCGATACGTGAGATTACCC | Closest flanking marker for *Hsf1* interval | INDEL marker |
| 410984-R | TGGCTGACTGTTCCACTACG | | |
| 391087-F | GTGTGAAACAGTGGCGAAGC | Closest flanking marker for *Hsf1* interval | Can be used as CAPS marker using enzyme *Acu*I |
| 391087-R | TAATGTATGCCCGAGAAACTTAGG | | |
| 17b-F | GCCACACTGAAGCACTCATA | *Hsf1-1603* genotyping assay | The *Hsf1* amplification fragment is 1345-bp and the wild type B73 fragment is 1010-bp |
| 18b-R | CAGCCGCAGCAACTCTGAGG | | |
| P3RCb-F | TGGATACGAGCTCCTCAAGAAGAT | *abph1* genotyping assay | The wild type B73 amplification fragment is 329-bp and the mutant *abph1* fragment is 430-bp. The P3RCb-F primer is in exon 3 and the Abph1-2-R primer is in exon 5. The AbP7b-R primer is specific for the retrotransposon insertion in the *abph1-R* allele. |
| Abph1-2-R | AATCCTCGGCGCCTTCCTCCA | | |
| Abp7b-R | CTGGTCTAGTGGACCCCACA | | |

**Supplemental Table 4.** Primers used for expression analysis

| Primer | Sequence (5' > 3') | Purpose | Gene ID |
|---|---|---|---|
| ABPH1-1 | AGGCGATGGCGAGCCGCAAG | *ZmRR3*; in situ; Fwd | GRMZM2G035688 |
| ABPH1-2 | AATCCTCGGCGCCTTCCTCCA | *ZmRR3*; in situ; Rev | |
| RR3-F2 | AGGATTTCCTGCTGAAGC | *ZmRR3*; RT PCR; Fwd | |
| RR3-R1 | GACACAGAGCTTCGGAAT | *ZmRR3*; RT PCR; Rev | |
| ARV0111 | GTGCCGTCCTATACTCGATCCG | *ZmRR2;* RT PCR; Fwd | GRMZM2G392101 |
| ARV0112 | TATACACAGGTGCAGTGCAGGG | *ZmRR2;* RT PCR; Rev | |
| ARV0119 | TCACCATCTTGCCTCTGTCCAG | *ZmHK1;* RT PCR; Fwd | GRMZM2G151223 |
| ARV0120 | TTGTGAGAGGCCATGTCAGAGG | *ZmHK1;* RT PCR; Rev | |
| HK1-A1 | AGAAGAACGGTCAGTTGTCG | *ZmHK1*; RT-PCR; Fwd | |
| HK1-A2 | CAACTCCATTTGCCTTATTTGTCC | *ZmHK1*: RT-PCR; Rev | |
| ARV0115 | CATGAACAAGCACAGGTGGGAC | *ZmCKO4b*; RT PCR; Fwd | GRMZM2G024476 |
| ARV0116 | CCACCAGGTAGAACACGTCCTC | *ZmCKO4b*; RT PCR; Rev | |
| ARV0079 | ATCTCGTTGGGGATGTCTTG | FPGS qPCR control, Fwd | GRMZM2G393334 |
| ARV0080 | AGCACCGTTCAAATGTCTCC | FPGS qPCR control, Rev | |
